# Systematic analysis of SARS-CoV-2 infection of an ACE2-negative human airway cell

**DOI:** 10.1101/2021.03.01.433431

**Authors:** Maritza Puray-Chavez, Kyle M. LaPak, Travis P. Schrank, Jennifer L. Elliott, Dhaval P. Bhatt, Megan J. Agajanian, Ria Jasuja, Dana Q. Lawson, Keanu Davis, Paul W. Rothlauf, Heejoon Jo, Nakyung Lee, Kasyap Tenneti, Jenna E. Eschbach, Christian Shema Mugisha, Hung R. Vuong, Adam L. Bailey, D. Neil Hayes, Sean P.J. Whelan, Amjad Horani, Steven L. Brody, Dennis Goldfarb, M. Ben Major, Sebla B. Kutluay

## Abstract

Established *in vitro* models for SARS-CoV-2 infection are limited and include cell lines of non-human origin and those engineered to overexpress ACE2, the cognate host cell receptor. We identified human H522 lung adenocarcinoma cells as naturally permissive to SARS-CoV-2 infection despite complete absence of ACE2. Infection of H522 cells required the SARS-CoV-2 spike protein, though in contrast to ACE2-dependent models, spike alone was not sufficient for H522 infection. Temporally resolved transcriptomic and proteomic profiling revealed alterations in cell cycle and the antiviral host cell response, including MDA5-dependent activation of type-I interferon signaling. Focused chemical screens point to important roles for clathrin-mediated endocytosis and endosomal cathepsins in SARS-CoV-2 infection of H522 cells. These findings imply the utilization of an alternative SARS-CoV-2 host cell receptor which may impact tropism of SARS-CoV-2 and consequently human disease pathogenesis.

## INTRODUCTION

Severe acute respiratory syndrome coronavirus 2 (SARS-CoV-2) emerged in late 2019 as the causative agent of the ongoing Coronavirus Disease-2019 (COVID-19) pandemic (Wu et al., 2020; Zhou et al., 2020). SARS-CoV-2 is a β-coronavirus and belongs to the larger group of coronaviruses (CoV) characterized by single-stranded, positive-sense RNA genomes of unusually large size (27-32 kb). Severe COVID-19 is marked by virus-induced lung damage (Wu and McGoogan, 2020), elevated levels of pro-inflammatory cytokines, immune cell infiltration in the lung (Chen et al., 2020; Huang et al., 2020) and multi-system involvement (Varga et al., 2020). The emergence of new SARS-CoV-2 variants bearing mutations in the viral spike (S) protein and recent reports of alternative viral entry mechanisms (Cantuti-Castelvetri et al., 2020; Clausen et al., 2020; Daly et al., 2020; Wang et al., 2021) demands comprehensive understanding of viral entry, replication and the host cell response. This knowledge will empower new therapeutics and vaccines to thwart future viral outbreaks.

SARS-CoV-2 homotrimeric viral S protein binding to the host cell angiotensin-converting enzyme 2 (ACE2) receptor mediates viral entry (Hoffmann et al., 2020; Letko et al., 2020; Walls et al., 2020; Zhou et al., 2020). High-sensitivity RNA *in situ* mapping revealed the presence of *ACE2* throughout the respiratory tract with highest expression in the nasal epithelium and gradually decreasing expression throughout the lower respiratory tract (Hou et al., 2020). Though present, ACE2 expression is relatively low in the respiratory tract (Aguiar et al., 2020; Hikmet et al., 2020) compared with higher levels in the gastrointestinal tract, kidney and myocardium (Hamming et al., 2004; Qi et al., 2020; Sungnak et al., 2020; To and Lo, 2004; Zhao et al., 2020; Zou et al., 2020). Low levels of ACE2 expression may be compensated by additional attachment/entry factors that enhance viral entry. For example, recent studies revealed that neuropilin-1 (NRP1) and heparan sulfate can facilitate ACE2-dependent SARS-CoV-2 entry *in vitro* (Cantuti-Castelvetri et al., 2020; Clausen et al., 2020; Daly et al., 2020). Additionally, the tyrosine-protein kinase receptor AXL mediates SARS-CoV-2 S pseudotyped lentivirus entry in an ACE2-independent manner, though the impact of AXL on the entry and the replication of fully infectious SARS-CoV-2 entry was significantly lower (Wang et al., 2021).

Multiple cell lines are routinely used to study β-coronavirus infection. SARS-CoV-2 primarily infects ciliated and type 2 pneumocyte cells in the human lung (Schaefer et al., 2020). As such, differentiated primary airway epithelial cells likely represent the most physiologically relevant model to study SARS-CoV-2 infection in culture. However, these cells require culture at an air-liquid interface, complex media formulations, and weeks of differentiation. Genetic manipulation, culture scalability, and donor-to-donor variability further complicate their use. Vero cells are derived from African green monkey kidney and are commonly used to propagate and study SARS-CoV-2 (Cagno, 2020; Chu et al., 2020b). The exceptional permissiveness of Vero cells is likely due to an ablated type I interferon response (IFN) due to a large deletion in the type I IFN gene cluster (Desmyter et al., 1968; Diaz et al., 1988; Osada et al., 2014). The inactivation of the type I IFN response and the presence of species-specific responses to viral pathogens (Long et al., 2019; Malim and Bieniasz, 2012; Rothenburg and Brennan, 2020) limits the utility and physiological relevance of Vero cell infection experiments. Although human Caco-2 colorectal adenocarcinoma and Huh-7 hepatocellular carcinoma cell lines support SARS-CoV-2 replication in an ACE2-dependent manner (Chu et al., 2020b; Kim et al., 2020; Ou et al., 2020), the only human lung cell line reported to be permissive to SARS-CoV-2 replication is Calu-3, albeit with significantly lower replication compared to Vero cells (Chu et al., 2020b; Ou et al., 2020). The general lack of permissiveness to SARS-CoV-2 infection in lung-derived cell lines is rescued by ectopic overexpression of the ACE2 receptor, suggesting that viral entry constitutes a major block to virus replication (Blanco-Melo et al., 2020).

In addition to ACE2 expression, cellular tropism of SARS-CoV-2 may be determined by cell intrinsic and innate immune defenses. Recognition of viral replication intermediates, typically viral nucleic acids, by toll-like receptors (TLRs) and RIG-I-like receptors (RLRs) culminate in secretion of type I and type III IFNs (Lazear et al., 2019; Schoggins, 2018). Type I/III IFN signaling results in the upregulation of numerous interferon-stimulated genes (ISGs) which collectively establish an antiviral state. SARS-CoV-2 induces lower levels of type I/III IFNs *in vitro* compared with other respiratory pathogens (Blanco-Melo et al., 2020; Chu et al., 2020a; Stukalov et al., 2020), possibly due to the expression of the SARS-CoV-2 proteins Nsp1 and ORF6 (Xia et al., 2020). Type I IFN pretreatment of cell culture models potently suppresses SARS-CoV-2 replication (Lokugamage et al., 2020; Xie et al., 2020), suggesting a potentially important role of this pathway in defining cellular tropism.

To identify new lung and upper airway cell culture models that are naturally permissive to SARS-CoV-2 infection, we infected a panel of human lung and head/neck cancer cell lines expressing varying levels of ACE2 and the TMPRSS2 protease. Unexpectedly, we found that the H522 lung adenocarcinoma cell line, which does not express any detectable levels of ACE2 or TMPRSS2, supports efficient SARS-CoV-2 replication. Infection of H522 cells is independent of ACE2, requires the viral S protein, and is suppressed by impeding clathrin-mediated endocytosis (CME) or endosomal cathepsin function. Time-resolved transcriptomic and proteomic profiling of infected H522 cells identified a robust activation of type I IFN responses in a MDA5-dependent manner, activation of CME, and modulation of cell cycle associated genes and pathways. Chemical inhibition of the AAK1 kinase, which potentiates CME, suppressed SARS-CoV-2 infection of H522 cells as well as primary human airway cultures. The ACE2 and TMPRSS2-independent infection of H522 cells establishes the presence of an alternative entry pathway for SARS-CoV-2 in human airway cells. Comprehensive understanding of these entry mechanisms may better explain the complex Covid-19 disease pathogenesis and the design of new and effective therapies.

## RESULTS

### H522 human lung adenocarcinoma cell line is permissive to SARS-CoV-2 infection

To identify additional cell types to model SARS-CoV-2 infection, we performed RNA-seq analysis on a panel of 10 lung and head/neck cancer cell lines, revealing varied expression levels of *ACE2* and *TMPRSS2* as well as other entry factors including proteases involved in S cleavage (*FURIN, CTSB, CTSL*) and neuropilin-1 (*NRP1*) (Cantuti-Castelvetri et al., 2020; Daly et al., 2020) (**Fig. 1A, Fig. S1 and Table S1**). Normalized RNA-seq read counts for established SARS-CoV-2 cell models Caco-2, Calu-3 and Vero E6 enabled comparative analysis (**Fig. 1A, Fig. S1 and Table S1**). Validation by qRT-PCR and protein quantification by immunoblotting showed ACE2 protein levels ranging from undetected to 2-3-fold lower than Vero E6 cells, currently the most permissive cell model to SARS-CoV-2 infection (**Fig. 1B, C**). The observed cell line-dependent differences in ACE2 protein migration is possibly due to post-translational modifications, including glycosylation (**Fig. 1C**) (Shajahan et al., 2020; Walls et al., 2020; Wrapp et al., 2020; Yang et al., 2020). Each cell line was then infected with SARS-CoV-2 before quantification of cell-associated viral RNA at 4- and 72-hours post infection (hpi) (**Fig. 1D**). The majority of cell lines, including those that express ACE2 and TMPRSS2 at relatively high levels (i.e. Detroit562 and H596) were not permissive to SARS-CoV-2 replication (**Fig. 1D**). H522 and to a lesser degree HCC827 cells supported virus replication (**Fig. 1D**). Surprisingly, neither ACE2 nor TMPRSS2 were detected in H522 cells (**Fig. 1A-D**).

**Figure 1.**
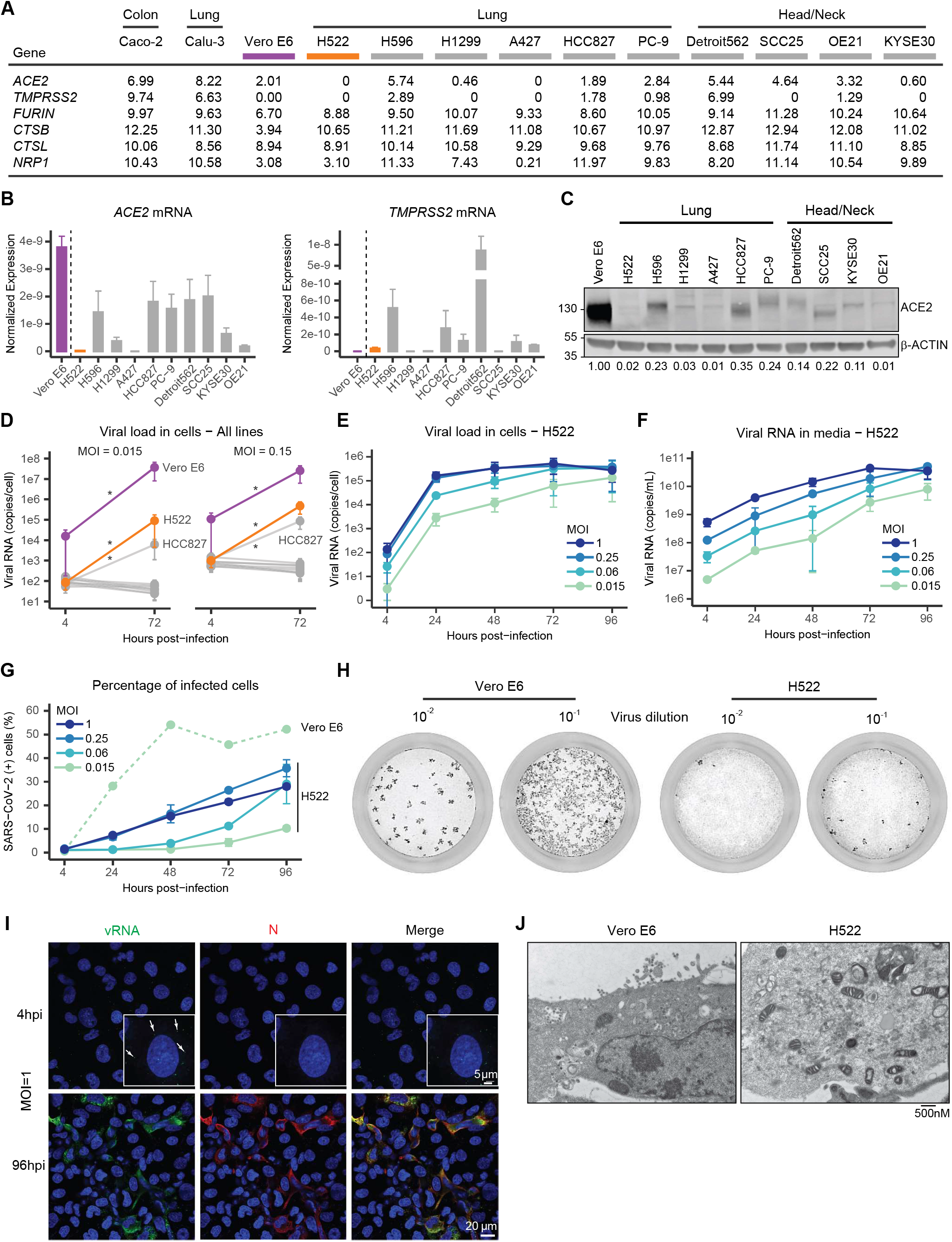
The H522 cell line is null for ACE2 expression and is permissive to SARS-CoV-2 infection. **A,** Normalized RNA-seq reads were aligned to the GRCh38 and Vervet-African green monkey genomes and quantified with Salmon (v1.3.0). The read counts for *ACE2, TMPRSS2, FURIN, CTSB, CTSL*, and *NRP1* are given for the indicated cell lines. See also Figure S1 and Table S1. **B,** qRT-PCR for *ACE2* and *TMPRSS2* expression normalized to 1μg input RNA for each cell line. *Cercopithecus aethiops* specific primers against *TMPRSS2* were used for the Vero E6 samples. Each bar represents mean, error bars indicate SEM (n=3). **C,** Immunoblot showing ACE2 expression across 10 lung and upper airway cancer cell lines and Vero E6 cells (representative of n=3). ACE2 expression was quantified using Licor Image Studio software in which ACE2 levels were normalized to β-ACTIN, set relative to Vero E6, and are indicated below the immunoblots. **D,** qRT-PCR for cell-associated SARS-CoV-2 RNA at 4 and 72 hpi at MOI=0.015 or 0.15. MOIs were determined by titration on Vero E6 cells. Error bars represent SEM (n=3). * indicates p<0.05 where significance was determined using two-way ANOVA and the Šidák correction for multiple comparisons. **E,** qRT-PCR for cell-associated SARS-CoV-2 RNA in H522 cells across various time points and MOIs. Error bars represent SEM (n=2). **F,** qRT-PCR for SARS-CoV-2 RNA in the supernatant of H522 cells across various time points and MOIs. Error bars represent SEM (n=2). **G,** Percent of SARS-CoV-2 infected H522 and Vero E6 cells determined by FACS for Nucleocapsid positive cells across various time points and MOIs. Error bars represent SEM (n=2). **H,** Plaque assays on H522 and Vero cells using two viral dilutions (10^−2^ and 10^−1^). Data are representative of three independent experiments. **I**, Representative images of H522 cells infected with SARS-CoV-2 at MOI=1. H522 cells were fixed and stained for SARS-CoV-2 RNA (green) by RNAScope reagents and Nucleocapsid (N) protein (red) at 4 and 96hpi and imaged by confocal microscopy (representative of n=2). See also Figure S2. **J**, Representative images using transmission electron microscopy (TEM) on Vero E6 and H522 cells infected with SARS-CoV-2 (MOI=0.1 pfu/cell and 24 hpi for Vero, MOI: 1 pfu/cell and 96 hpi for H522).

Given the possibility of ACE2-independent infection, we focused our efforts on H522 cells. To define viral replication kinetics, H522 cells were infected at various multiplicities of infection (MOIs) before monitoring for virus growth over 4 days. Cell-associated viral RNA levels increased substantially (3-4 logs) within 24 hours of infection in a MOI-responsive manner (**Fig. 1E)**and corresponded to heightened viral RNA in the cell culture supernatants (**Fig. 1F**). We confirmed replication competency of virus released from H522 cells through plaque assays on Vero cells (up to 2.2×10^5^ pfu/mL, data not shown). While permissive, infection progressed slower in H522 cells compared to Vero E6 cells and higher doses of the virus were required to achieve similar numbers of infected cells (**Fig. 1G)**. Viruses formed plaques on H522 cells and plaque sizes were comparable to those obtained on Vero E6 cells, albeit the effective MOI was ~20-fold lower (**Fig. 1H**).

Quantified RNA-*in situ* hybridization (ISH) revealed the kinetics of SARS-CoV-2 viral replication and spread in H522 cells (**Fig. 1I, S2A, B**). Incoming virions were readily detected at 4 hpi by RNA-ISH in cells infected with a MOI of 1 (white arrows to green puncta; **Fig. 1I**). Expectedly, both the number of cells positive for viral RNA and the number of viral RNA puncta/cell were MOI-dependent (**Fig. S2A, B**). We tracked viral spread over time and observed increased staining for both viral RNA and N, and increased number of infected cells per field (**Fig. S2A**). Furthermore, similar to Vero cells, virions were observed in membrane-limited compartments in infected H522 cells, and frequently in apoptotic cells (**Fig. 1J**). These results show that H522 cells are productively infected with SARS-CoV-2 despite lacking any detectable levels of ACE2/TMPRSS2 expression.

### SARS-CoV-2 spike (S) protein is necessary but not sufficient for H522 infection

SARS-CoV-2 entry into host cells requires viral S-mediated engagement of the ACE2 receptor and priming of S by the TMPRSS2 or other host cell proteases (Hoffmann et al., 2020; Letko et al., 2020; Walls et al., 2020; Zhou et al., 2020). Given that ACE2 and TMPRSS2 mRNA and protein were undetectable in H522 cells (**Fig. 1A-C**), we tested the dependency on S for SARS-CoV-2 infection. H522 parental cells, H522 cells stably expressing ACE2 (H522-ACE2) and Vero E6 cells were infected with SARS-CoV-2 in the presence of a S neutralizing antibody (Alsoussi et al., 2020) or soluble human ACE2-Fc decoy receptor (**Fig. 2A-C**) (Case et al., 2020). In both experiments, blockage of S significantly diminished the amount of cell-associated viral RNA in H522, H522-ACE2 and Vero E6 cells (**Fig. 2B-C**). The decreased sensitivity of H522-ACE2 cells to treatment as compared to parental H522 cells likely reflects the high overexpression of ACE2 (**Fig. 2A**).

**Figure 2.**
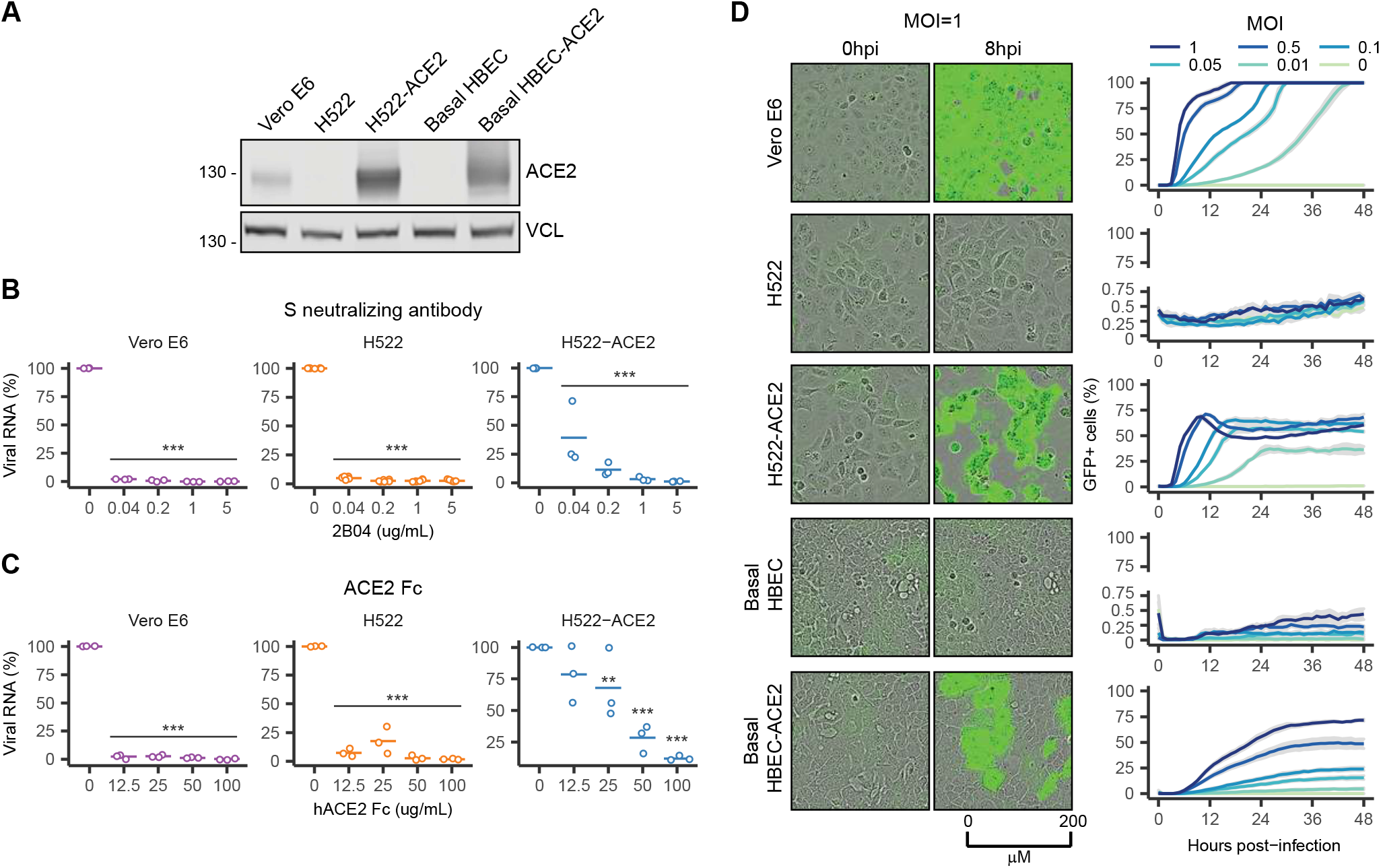
The SARS-CoV-2 S protein is necessary but not sufficient for viral entry in the H522 cell line. **A,** Representative immunoblot showing ACE2 expression and Vinculin as the loading control in Vero E6, H522, H522-ACE2, basal HBEC, and basal HBEC-ACE2 cells. **B**, Viruses were pre-treated with increasing concentrations of S neutralizing antibody for 1 h and then cells were infected with SARS-CoV-2 at MOI=0.1 in the presence of the S neutralizing antibody. Cell-associated SARS-CoV-2 RNA was detected by qRT-PCR at 24 hpi and was normalized to mock treated (n=3). *** indicates p<0.001 where significance was determined using two-way ANOVA and the Dunnett correction for multiple comparisons. **C,** SARS-CoV-2 viruses were pre-treated with increasing amounts of soluble ACE2-Fc for 1 h and then cells were infected with SARS-CoV-2 at MOI=0.1 in the presence of ACE2-Fc. Cell-associated SARS-CoV-2 RNA was detected by qRT-PCR at 24 hpi and was normalized to mock treated (n=3). ** indicates p<0.01 and *** indicates p<0.001 where significance was determined using two-way ANOVA and the Dunnett correction for multiple comparisons. **D,** Representative images of cells infected with VSV-SARS-CoV-2-S_Δ21_ at 0 and 8hpi using an Incucyte^®^ S3 Live Cell Analysis System (n=3). Percent GFP positive cells seeded in triplicate were quantified over time with the shaded grey region indicating standard deviation.

We next investigated whether S is sufficient for viral entry in H522 cells. For this, we used a replication-competent GFP-reporter vesicular stomatitis virus (VSV) that expresses a modified form of the SARS-CoV-2 S protein (designated VSV-GFP-SARS-CoV-2-S_Δ21_). The S protein present on the VSV particles is antigenically and functionally indistinguishable to the native S trimers in infectious SARS-CoV-2 (Case et al., 2020). We infected the following cell models with VSV-GFP-SARS-CoV-2-S_Δ21_: Vero E6, H522, H522-ACE2, primary basal human bronchial epithelial cells (HBECs) and HBECs engineered to express ACE2 (Basal HBEC-ACE2; **Fig. 2A**). While Vero E6, H522-ACE2 and basal-HBEC-ACE2 cells were readily infected, H522 and basal cells were resistant to infection by VSV-GFP-SARS-CoV-2-S_Δ21_ (**Fig. 2D**), suggesting that S protein is not sufficient for viral entry.

### SARS-CoV-2 replication in H522 cells is independent of ACE2

Two orthogonal approaches were used to test whether ACE2 mediated SARS-CoV-2 infection of H522 cells. First, cells were pre-treated with anti-ACE2 blocking antibody before addition of SARS-CoV-2. Anti-DC-SIGN and anti-GFP antibodies served as negative controls as well as the use of CHO-derived PgsA-745 cells, which are not permissive to infection (**Fig. 3A**). While the ACE2 blocking antibody significantly decreased the amount of cell-associated viral RNA in Calu-3 cells, it did not impact SARS-CoV-2 viral RNA levels in H522 cells (**Fig. 3A**). As expected, neither the anti-DC-SIGN nor the anti-GFP antibodies significantly affected viral RNA levels in H522 and Calu-3 cells, and virus replication remained at background levels in PgsA-745 cells.

**Figure 3.**
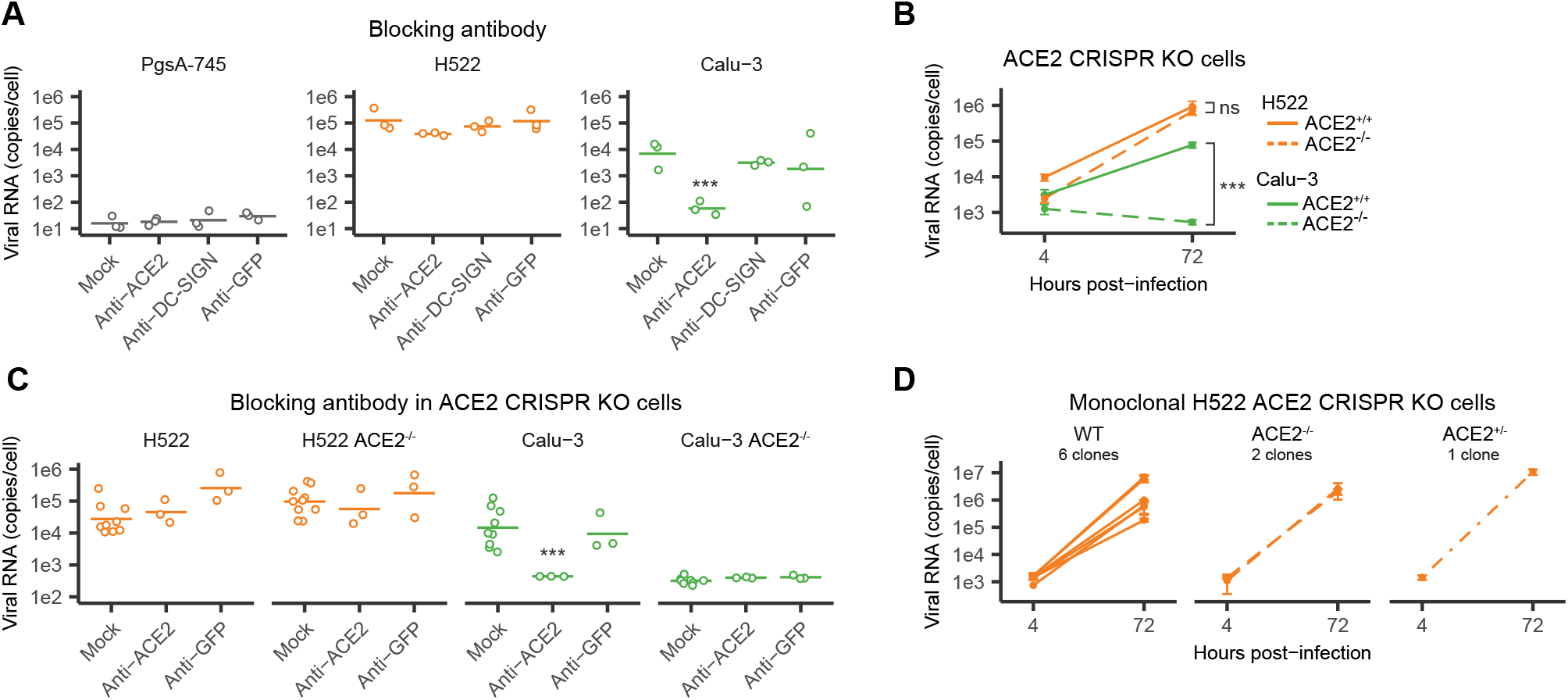
The H522 cell line is permissive to SARS-CoV-2 infection independent of ACE2 expression. **A**, Cells were pre-treated with 20 μg/ml of the indicated blocking antibodies for 1 h and then infected with SARS-CoV-2 at MOI=0.1 in the presence of the blocking antibodies. Cell-associated SARS-CoV-2 RNA was detected by qRT-PCR at 72 hpi (n=3). *** indicates p<0.001 where significance was determined using two-way ANOVA and the Dunnett correction for multiple comparisons. **B**, Polyclonal populations of H522 and Calu-3 (ACE2^+/+^ and ACE2^−/-^) cells were infected with SARS-CoV-2 virus and cell-associated SARS-CoV-2 RNA was detected by qRT-PCR 4 and 72 hpi (n=8). Error bars indicate the SEM. *** indicates p<0.001 where significance was determined using two-way ANOVA and the Tukey correction for multiple comparisons. See also Figure S3. **C**, Polyclonal populations of H522 and Calu-3 (ACE2^+/+^ and ACE2^−/-^) cells were pre-treated with 20 μg/ml of the indicated blocking antibodies for 1 h and then infected with SARS-CoV-2 at MOI=0.1 in the presence of the blocking antibodies. Cell-associated SARS-CoV-2 RNA was detected by qRT-PCR 72 hpi (n=3). Error bars indicate the SEM. *** indicates p<0.001 where significance was determined using two-way ANOVA and the Tukey correction for multiple comparisons. See also Figure S3. **D**, Monoclonal populations from H522 ACE2^+/+^ (6 clones), ACE2^−/-^ (2 clones), and ACE2^+/-^ (1 clone) were infected with SARS-CoV-2 at MOI=0.1 and cell-associated SARS-CoV-2 RNA was detected by qRT-PCR 4 and 72 hpi (n≥3). Error bars indicate the SEM. See also Figure S3.

In a second approach to test ACE2 involvement, we inactivated the ACE2 genetic locus by CRISPR gene editing in H522 and Calu-3 cells. Polyclonal cell populations containing CRISPR-edited loci were infected with SARS-CoV-2 and viral replication was monitored at 4 and 72 hpi (**Fig. 3B, S3A**). While viral RNA levels increased at similar levels in H522 and H522 ACE2^−/-^ cells, lack of ACE2 significantly reduced SARS-CoV-2 replication in Calu-3 cells (**Fig. 3B**). In agreement, the addition of an ACE2 blocking antibody did not impair virus replication in H522 or H522 ACE2^−/-^ cells, but completely abolished replication in Calu-3 cells (**Fig. 3C**). We next isolated monoclonal populations of H522 ACE2 WT (n=6), H522 ACE2^−/-^ (n=2) and H522 ACE2^+/-^ (n=1) cells to corroborate these findings (**Fig. S3B**). Sanger sequencing of the edited loci in two independent monoclonal populations revealed unique 5 bp deletions in Exon 3 of ACE2, resulting in the same truncated ACE2 protein lacking the C-terminal 672 amino acids, which includes the intracellular domain, transmembrane domain, collectrin domain and 75% of carboxypeptidase domain (**Fig. S3B**). SARS-CoV-2 infection of monoclonal cell lines from H522 control and H522 ACE2 KO resulted in similar levels of infection (**Fig. 3D**).

Taken together, these data establish that H522 cells are permissive to SARS-CoV-2 infection independent of ACE2 but dependent on the SARS-CoV-2 S protein.

### Clathrin-mediated endocytosis governs SARS-CoV-2 infection of H522 cells

To begin to decipher the mechanism(s) of SARS-CoV-2 entry into H522 cells, we performed infections in the presence of compounds that interfere with SARS-CoV-2 entry, including camostat mesylate (TMPRSS2 inhibitor) (Hoffmann et al., 2020; Shang et al., 2020; Shema Mugisha et al., 2020a), E64D (broad spectrum inhibitor of proteases, including endosomal cathepsins) (Ou et al., 2020; Shema Mugisha et al., 2020a), bafilomycin A (inhibitor of vATPase) (Ou et al., 2020; Shema Mugisha et al., 2020a) and apilimod (inhibitor of PIKfyve) (Kang et al., 2020; Ou et al., 2020; Shema Mugisha et al., 2020a). We additionally included a specific inhibitor of AAK1 kinase (SGC-AAK1-1); AAK1 promotes CME through phosphorylation of the AP2M1 subunit of the AP2 complex (Agajanian et al., 2019; Conner and Schmid, 2002, 2003). E64D, bafilomycin A, SGC-AAK1-1 and apilimod significantly reduced cell-associated viral RNAs in a dose-dependent manner, whereas camostat mesylate increased viral RNA levels (**Fig. 4A**). These findings were corroborated in comparative analysis of H522, Vero E6 and H522-ACE2 cells. Bafilomycin A significantly decreased cell-associated viral RNA levels in H522 and H522-ACE2 cells but did not affect viral entry in Vero E6 cells (**Fig. S4**). Inhibition of both AAK1 and endosomal cathepsins B/L significantly decreased viral RNA levels in H522 cells but did not impact ACE2-dependent replication at appreciable levels in Vero E6 and H522-ACE2 cells at this concentration (**Fig. S4**). While apilimod decreased viral entry in Vero E6 cells, the effect was modest in H522 cells and trended towards significance (p=0.07; **Fig. S4**). Finally, camostat mesylate did not decrease, and on the contrary, increased the amount of cell-associated viral RNA in H522s, highlighting the TMPRSS2 independence of viral entry (**Fig. S4**).

**Figure 4.**
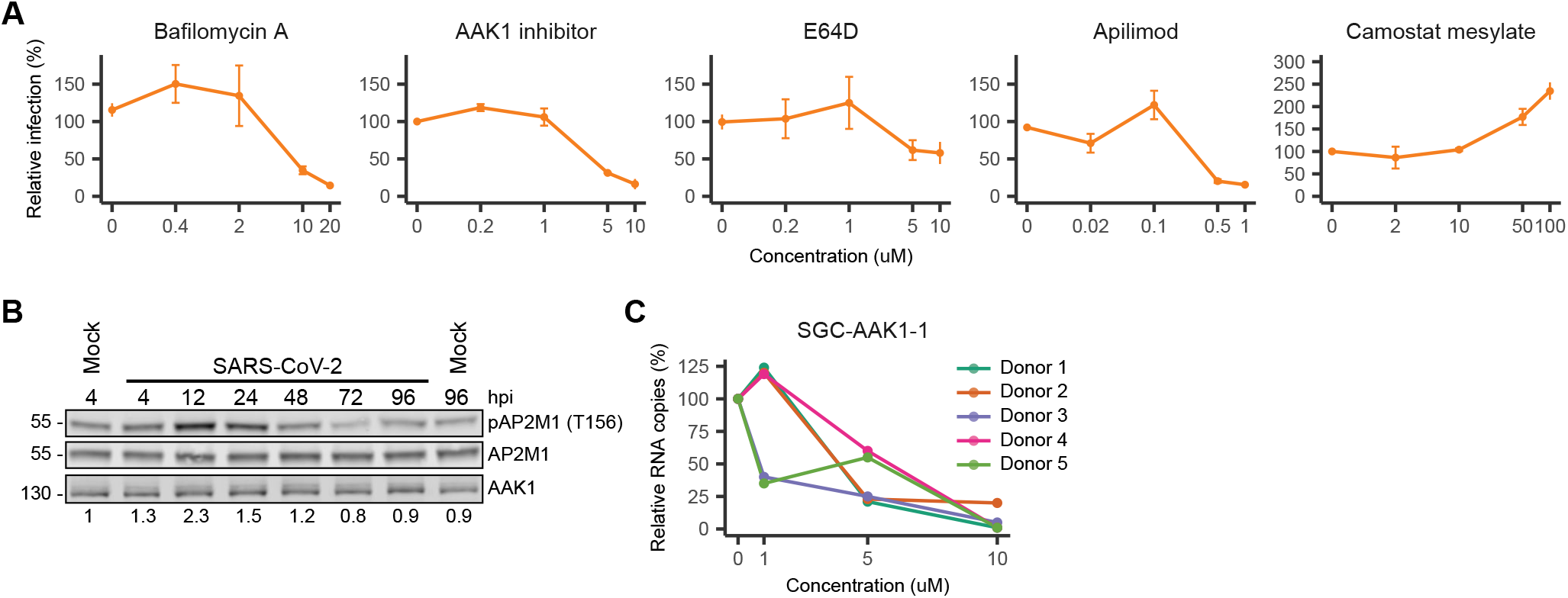
H522 infection by SARS-CoV-2 is dependent on clathrin-mediated endocytosis and endosomal cathepsins. **A**, H522 cells were pre-treated with increasing concentrations of bafilomycin A, SGC-AAK1-1, E64D, apilimod, or camostat mesylate for 1 h and then infected with SARS-CoV-2 at MOI=1 in the presence of the inhibitors. Cell-associated SARS-CoV-2 RNA was detected by qRT-PCR 24 hpi and normalized to DMSO treated cells (n≥3). See also Figure S4. **B**, Immunoblot showing pAP2M1 (T156), AP2M1, and AAK1 levels in H522 cells infected with SARS-CoV-2 over time (representative of n=2). pAP2M1 (T156) levels were normalized to total AP2M1 and set relative to the 4 hours mock control. Quantification was performed using the Licor Image Studio software and values are indicated below the immunoblots. **C**, Basal HBECs from 5 different donors were pre-treated with increasing concentrations of SGC-AAK1-1 for 2 h and then infected with SARS-CoV-2 in the presence of the inhibitor. Cell-associated SARS-CoV-2 RNA was detected by qRT-PCR 72 hpi and normalized to DMSO treated cells.

Western blot analysis of H522 cells infected with SARS-CoV-2 revealed transient induction of AP2M1 phosphorylation 12-24 hpi, further supporting the involvement of CME in H522 viral infections (**Fig. 4B**). AAK1 inhibitors are highly specific and have been considered to be viable therapeutic options for treatment of SARS-CoV-2 (Richardson et al., 2020). Consistent with our observations in H522 cells, inhibition of AAK1 kinase activity in differentiated primary HBECs grown at air-liquid interface led to a 10-20-fold decrease in cell-associated SARS-CoV-2 RNA in a dose responsive manner (**Fig. 4C**). Together, these data support a role for CME and endosomal cathepsins in SARS-CoV-2 infections of H522 cells.

### SARS-CoV-2-infected H522 cells demonstrate RNA-level upregulation of type I interferon responses and modulation of cell cycle genes

To determine how H522 cells respond to SARS-CoV-2 infection, we conducted RNA-seq on cells infected at high and low MOI and followed the infection over the course of 4 days (**Fig. 5A**). As expected, SARS-CoV-2 mRNA levels increased with time and MOI, plateauing around 24-48 hpi (**Fig. 5B**). At the peak of infection, 5-10% of total reads mapped to SARS-CoV-2 RNAs. Principal component analysis (PCA) showed samples separated well based on MOI and time post-infection (**Fig. 5C**). Analysis of differentially expressed genes (DEGs) at 96 hpi revealed MOI-dependent upregulation of IRF9, as well as numerous IFN-stimulated genes including ISG15, MX1, IFI35 and OAS3 (**Fig. 5D, Table S2**). Hierarchical consensus clustering of the 2,631 DEGs (|logFC|>2 and q<0.005) generated 7 temporally resolved clusters (**Fig. 5E, F**). Over-representation analysis of each cluster revealed an initial sharp increase of cell cycle regulatory and inflammatory genes followed by decreasing levels as the infection proceeded (cluster 1) (**Fig. 5F, G, Table S3**). Additionally, IFN-alpha/beta signaling and downstream ISGs significantly increased as early as 48 hpi and continued to increase further by 96 hpi, consistent with high levels of SARS-CoV-2 infection (cluster 4; **Fig 5E-G, Table S3**). Modulation of the cell cycle and sustained IFN signaling throughout infection were confirmed by gene set enrichment analysis (GSEA) of genes at each time point (**Fig. 5H, Table S3**). Together, these findings highlight global changes in the H522 transcriptome in response to SARS-CoV-2 and marked induction of antiviral immune responses.

**Figure 5.**
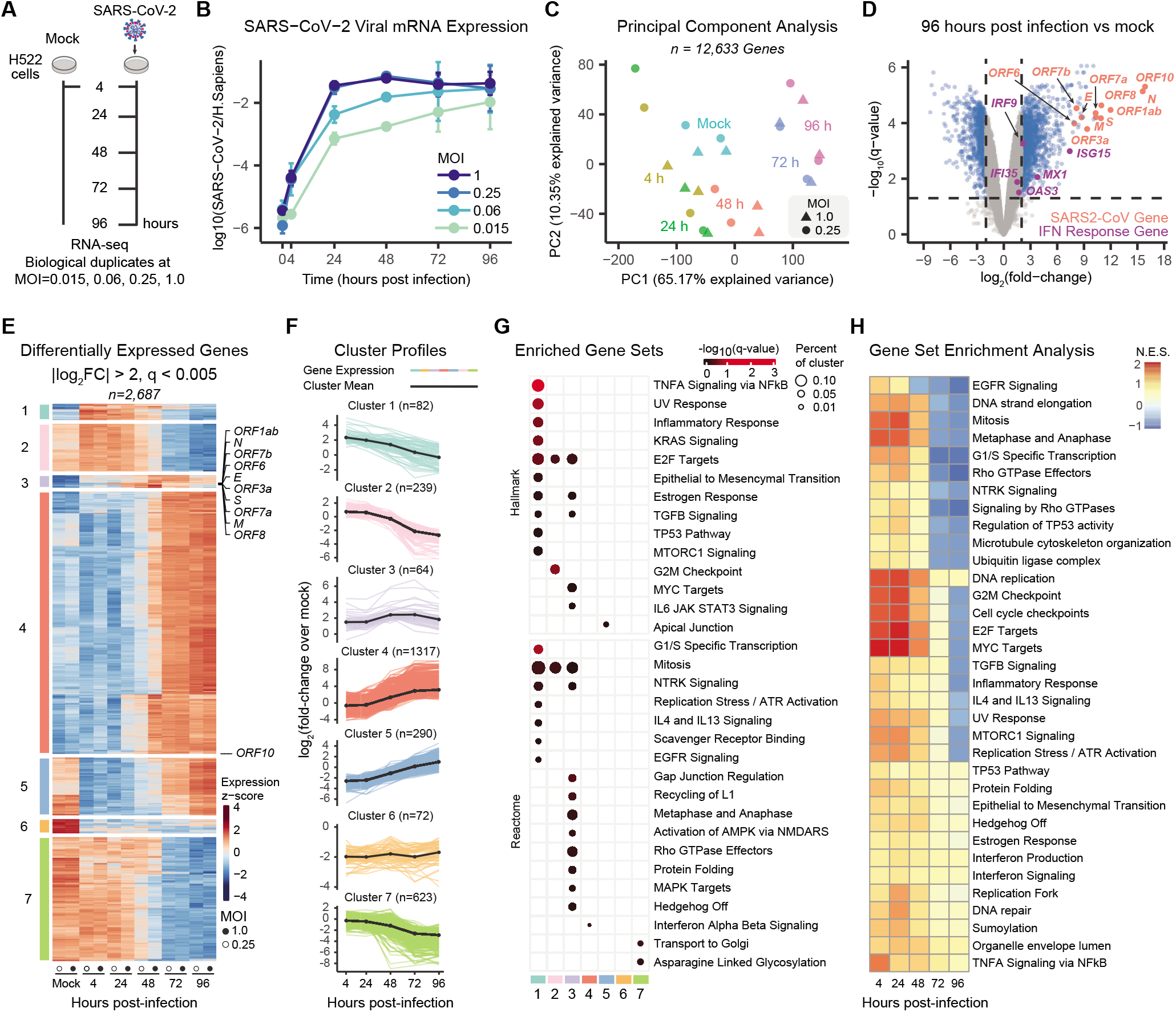
H522 transcriptome response to SARS-CoV-2 infection. **A**, Experimental design of transcriptomics experiments. H522 cells were infected with SARS-CoV-2 at MOI 1.0, 0.25, 0.06, or 0.015 and harvested after 4, 24, 48, 72, and 96 h. Mock-infected cells were harvested after 4 h. All conditions were performed in duplicate. **B**, Relative expression of SARS-CoV-2 RNA vs. *H. sapiens* RNA from H522 (n=2). **C**, Principle component analysis of highly expressed genes from MOIs 0.25 and 1 across all time points. **D**, Volcano plot of gene expression changes comparing mock infection to 96 hours post infection of MOIs=0.25 and 1. Select changes in IFN response genes (purple) and SARS-CoV-2 genes (salmon) are highlighted. See also Table S2. **E**, Hierarchical clustering of differentially expressed genes (DEGs) after infection. Genes were filtered for an absolute log_2_ fold change >2 and adjusted p-value < 0.005 at any time point. **F**, Log_2_ fold changes of DEGs as grouped by clustering. The colored lines represent quantification of an individual gene whereas the solid black represents the cluster mean. **G**, Hypergeometric enrichment analysis of biological gene sets in the identified gene clusters (D-E). See also Table S3. **H**, Rank-based gene set enrichment analysis. Gene sets were queried if identified by hypergeometric analysis RNA seq (5F) or proteomics data (6E). Display indicated p-adjusted < 0.05. N.E.S. = normalized enrichment score.

### SARS-CoV-2-induced proteome changes in H522 cells reveal induction of type I IFN, cell cycle and DNA replication pathways

To define the impact of SARS-CoV-2 infection on the H522 proteome, we conducted whole cell quantitative proteomics experiments over the course of 4 days (**Fig. 6A**). Biological triplicates for each time point were processed and the abundance of 7,469 proteins was analyzed across samples. PCA highlights the high level of reproducibility and clustering of samples by infection and time post-infection (**Fig. 6B**). Similar to viral RNAs, abundance of viral proteins increased substantially within the first 24 hours of infection and plateaued thereafter (**Fig. 6C**). At 96hpi vs 96h mock, 492 differentially regulated proteins were identified (**Fig. 6D**). Unsupervised clustering defined seven unique clusters that characterize the temporal regulation of the H522 proteome (**Fig. 6E, Table S4**). Overall, the majority of the differentially expressed proteins increased following SARS-CoV-2 infection, with proteins in cluster 4 displaying the greatest fold changes (**Fig. 6F, Table S4**). Over-representation analysis revealed that cluster 4 proteins include those involved in the IFN-α and IFN-γ responses, which were the most significantly altered pathways (**Fig. 6G, Table S5**). Of note, all viral proteins were present in Cluster 2 and their accumulation preceded the induction of type I/III IFNs (**Fig. 6E**). Cell cycle regulators were increased at early time points but declined thereafter, matching what was seen at the RNA level with a 12-24 hour delay (Cluster 1; **Fig.6 E-G**). Clusters 2 and 3 included proteins similarly involved in cell cycle regulation, DNA replication/repair, and microtubule organization but tended to remain upregulated during SARS-CoV-2 infection (**Fig. 6E-G**). Finally, clusters 6 and 7 included proteins that were downregulated and included plasma membrane proteins such as Semaphorins (SEMA3A, C, D), APOE, ERBB4, LRP1, and SLIT2 with potential roles in viral entry pathways (**Fig. 6E-G**). We next looked for genes that correlated between our transcriptomic and proteomic datasets. While we see an even distribution of correlations when including all genes, there is an increase in correlated genes when focusing on only the differently expressed proteins (**Fig. 6H**). Among these genes, only the IFN-α and IFN-γ signaling pathways were identified by GSEA for enrichment in correlation, further supporting an IFN response in H522 cells to SARS-CoV-2 infection (**Fig. 6I**).

**Figure 6.**
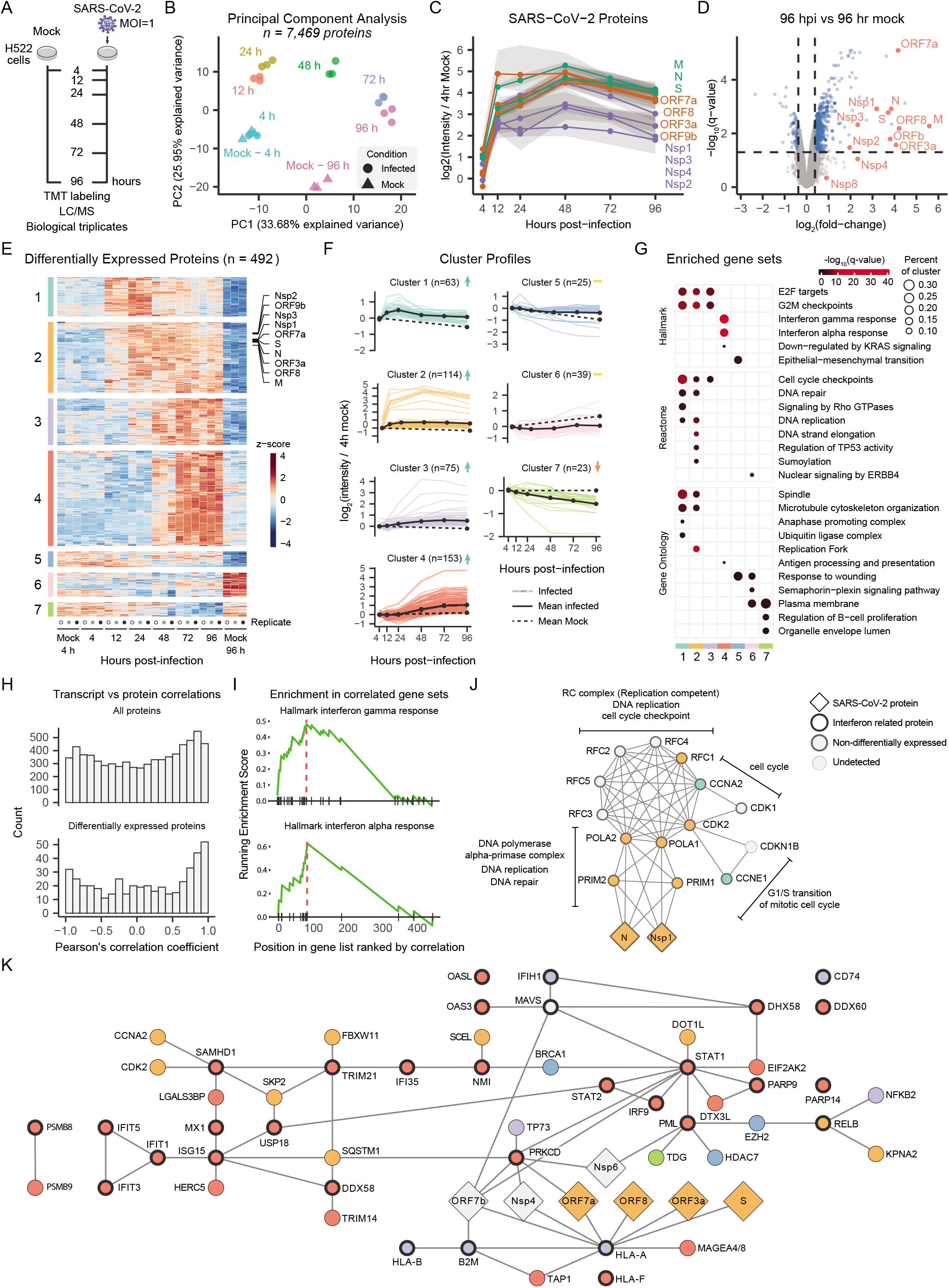
H522 infection with SARS-CoV-2 results in proteome changes within the type I IFN, cell cycle, and DNA replication pathways. **A**, Experimental design of proteomics experiments. H522 cells were infected with SARS-CoV-2 at MOI=1 and harvested after 4, 12, 24, 48, 72, and 96 h. Mock-infected cells were harvested after 4 and 96 h. Peptides labeled with TMT10 reagents were analyzed by liquid chromatography-mass spectrometry. **B**, Principal component analysis of whole cell proteomics of H522 cells infected with SARS-CoV-2 across a 4-day time course (n=3). **C,** Quantification of total ion intensities for each identified SARS-CoV-2 protein over time and normalized to the 4 h mock control. The shaded grey regions represent SEM. **D**, Volcano plot of protein abundance at 96hpi compared to the 96 h mock control. See also Table S4 **E**, Differentially expressed proteins from ‘D’ were clustered based on z-score. **F**, Quantification of total ion intensities normalized to the 4 h mock control for each protein across the 7 identified clusters in ‘D’. The colored lines represent quantification of an individual protein whereas the solid black and dashed black lines represent the mean of infected and mock samples, respectively. **G**, Hypergeometric enrichment analysis from three different databases for each individual cluster in ‘D’ (Hallmark, Reactome, Gene Ontology). The color of the circle represents significance (q-value), whereas the size of the circle indicates the percentage of the cluster represented in the pathway. See also Table S5. **H**, Distribution of Pearson’s correlation coefficient between a gene’s transcript and protein log_2_ fold change over 4 h mock for all proteins and differentially expressed proteins. Correlations used the matching time points of 4, 24, 48, 72, 96 hpi. **I**, Rank-based gene set enrichment analysis. Differentially expressed proteins were ranked by their correlation to transcript levels. **J**, Protein complexes of differentially expressed H522 and SARS-CoV-2 proteins associated with DNA replication and cell cycle checkpoint. Complexes and functions were extracted from the CORUM database. The colors correspond to the whole cell proteomic clusters identified in ‘D’. See also Figure S5. **K**, Protein interaction network of differentially expressed H522 and SARS-CoV-2 proteins associated with the IFN response. Interactions were determined from the BioGRID Multi-Validated Datasets. Interferon related functions were extracted from GO terms in MSigDB. The colors correspond to the whole cell proteomic clusters identified in ‘D’. See also Figure S5.

To further illuminate pathways altered by SARS-CoV-2 infection, we mined the CORUM database for protein complexes consisting mostly of differentially expressed proteins (**Fig. 6J, S5**). In total, 27 complexes were found and involved IFN signaling, cell cycle/DNA replication, DNA repair, epigenetic modification, and protein folding/ubiquitination, (**Fig. 6J-K, S5**). Over half of the complexes had functions in cell cycle and DNA repair (n=18). Of note, the viral proteins N and Nsp1 were previously reported to interact with components of the DNA polymerase alpha-primase complex, suggesting that the observed protein level changes are a direct result of these interactions (**Fig. 6J**). Additionally, we generated protein interactions networks based on BioGRID Multi-Validated Datasets for the 492 differentially expressed proteins in H522 cells and the SARS-CoV-2 viral proteome (**Fig. 6K**). A network emerged that contained 53 proteins involved in IFN signaling and downstream ISGs (**Fig. 6K**). Seven viral proteins associate with various host proteins within this network, raising the possibility that SARS-CoV-2 may directly modulate the IFN response in H522 cells (**Fig. 6K**). Taken together, these results show that SARS-CoV-2 infection of H522 cells leads to significant upregulation of several genes involved in innate immune pathways and cell cycle regulation at both the mRNA and protein level.

### MDA5 mediates the sensing of SARS-CoV-2 replication intermediates

Transcriptional profiling and proteomics revealed the IFN signaling pathway as the major immune signaling pathway responding to SARS-CoV-2 infection in H522 cells (**Fig. 5 and 6**). To validate the IFN response, we measured the levels and activation of STAT1 and downstream ISGs in infected H522 cells. We found that SARS-CoV-2 replication induced both upregulation of STAT1 expression and its phosphorylation, as well as downstream ISGs MX1 and IFIT1 by 48 hpi (**Fig. 7A**). MX1 and IFIT1 were upregulated further as the infection progressed at 72 and 96 hpi (**Fig. 7A**). Upregulation of the type I IFN response was delayed relative to the accumulation of viral N protein expression which peaked by 24 hpi, possibly due to antagonism of host responses at early times in infection.

**Figure 7.**
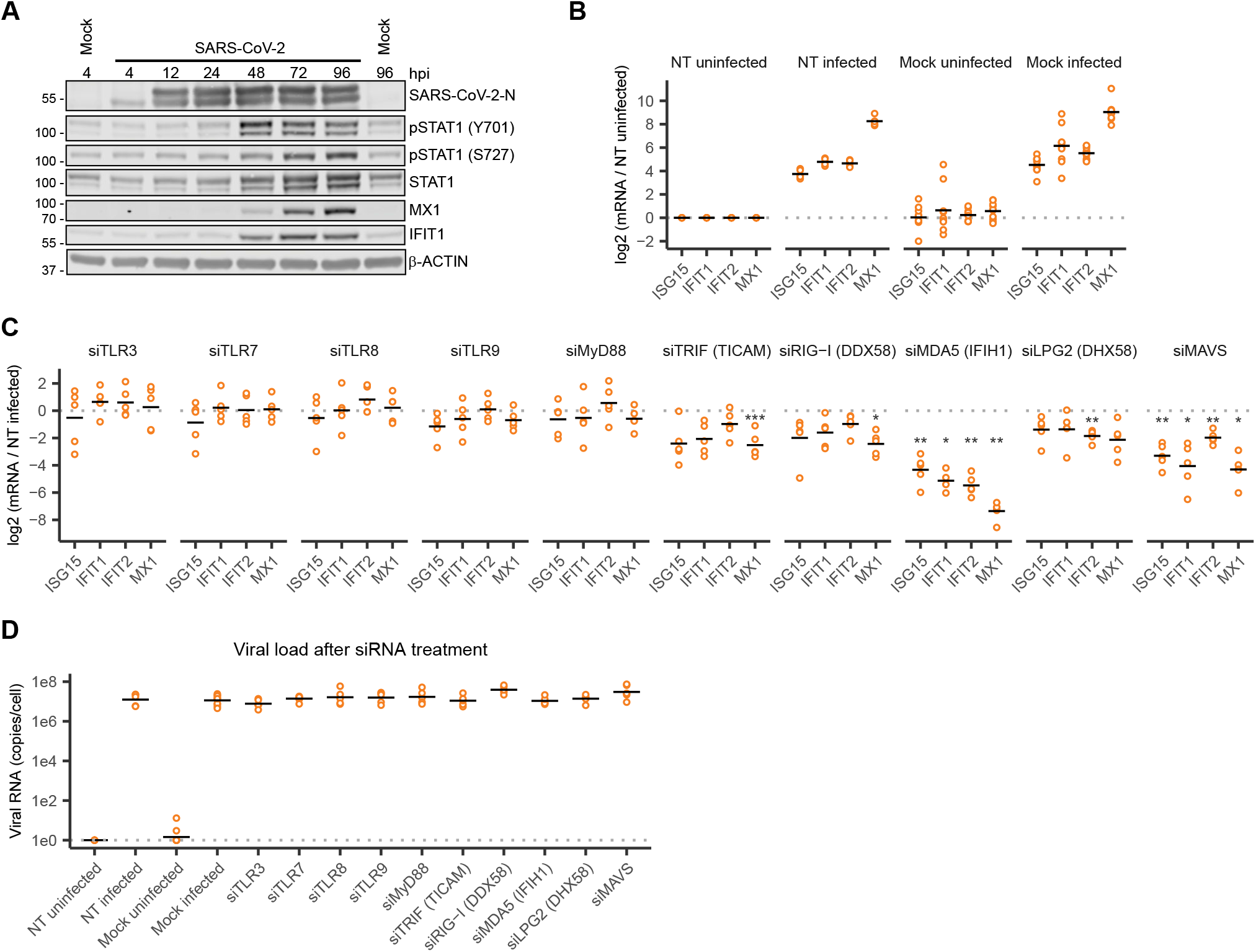
MDA5 mediates the IFN response to SARS-CoV-2 infection. **A**, Immunoblot depicting the IFN response in H522 cells infected with SARS-CoV-2 over time (representative of n=2). β-actin represents the loading control. **B**, ISG mRNA levels was detected by qRT-PCR in H522 cells infected with SARS-CoV-2 96 hpi. H522 cells were either mock transfected or transfected with a non-targeting (NT) siRNA 24 h prior to infection. **C**, ISG mRNA levels was detected by qRT-PCR in H522 cells infected with SARS-CoV-2 96 hpi. H522 cells were transfected with a non-targeting (NT) siRNA or a panel of siRNAs targeting genes involved in viral sensing 24 h prior to infection. * indicates p<0.05, ** indicates p<0.01 and *** indicates p<0.001 where significance was determined using two-way ANOVA and the Dunnett correction for multiple comparisons. See also Figure S6. **D**, qRT-PCR for cell-associated SARS-CoV-2 RNA in H522 cells 96hpi. H522 cells were transfected with a non-targeting (NT) siRNA or a panel of siRNAs targeting genes involved in viral sensing 24 hours prior to infection.

We next sought to define the mechanism by which H522 cells sense and respond to SARS-CoV-2 replication. Components of TLR and RLR-dependent sensing pathways were knocked down by siRNA transfection and upregulation of ISGs assessed following SARS-CoV-2 infection. Most targets remained efficiently knocked down up to 120 hours post transfection (**Fig. S6**). The low knockdown efficiency of TLR3, TLR7, TLR8 and TLR9 is most likely due to their low to undetectable basal expression levels (**Fig S6**). In line with RNA-seq and proteomics findings, ISG15, IFIT1, IFIT2 and MX1 were upregulated upon infection in mock- and non-targeting (NT) siRNA-transfected H522 cells (**Fig. 7B**). Depletion of MDA5 and, to a lesser extent, the downstream adaptor MAVS significantly reduced ISG induction in response to SARS-CoV-2 infection (**Fig. 7C)**. Knockdown of all the other targets had little or no impact on ISG upregulation (**Fig. 7C)**. Despite the decreased IFN response in MDA5 and MAVS depleted H522 cells, SARS-CoV-2 viral RNA levels remained at similar levels compared to controls (**Fig. 7D**). These results together suggest that viral RNAs are sensed by components of the RLR pathway in H522s resulting in activation of the type I IFN response.

## DISCUSSION

Our screen of human lung and head/neck cancer cell lines that express varying levels of ACE2 and TMPRSS2 identified the H522 human lung adenocarcinoma cell line as being naturally permissive to SARS-CoV-2 infection despite no evidence of ACE2 and TMPRSS2 expression. Using CRISPR editing and neutralizing antibodies, we confirmed the ACE2 independence of H522 infection, a paradigm shifting finding which suggests the utilization of an alternative receptor in a cell line of lung origin. As ACE2 expression is comparably low in the human respiratory system (Aguiar et al., 2020; Hikmet et al., 2020), other co-receptors and/or attachment factors have been suspected to enhance viral entry. Indeed, recent findings establish NRP1 and heparan sulfate as positive mediators of ACE2-dependent SARS-CoV-2 entry (Cantuti-Castelvetri et al., 2020; Clausen et al., 2020; Daly et al., 2020). Given their dependency on ACE2 expression, it is unlikely that these factors mediate ACE2-independent entry into H522 cells. In addition, coronaviruses can utilize a diverse array of glycoconjugates as attachment factors. For example, human coronaviruses OC43 (HCoV-OC43) and HKU1 (HCoV-HKU1) bind to 9-*O*-Ac-sialosides (Huang et al., 2015; Hulswit et al., 2019; Tortorici et al., 2019), and MERS-CoV (a β-coronavirus) as well as α- and γ-coronaviruses bind to other sialoglycans distinct from 9-*O*-Ac-sialosides (Li et al., 2017; Liu et al., 2015; Park et al., 2019; Schultze et al., 1996; Vlasak et al., 1988; Wickramasinghe et al., 2011). Whether sialic acids serve as entry receptors and/or cell attachment factors for SARS-CoV-2 in H522 cells remains to be determined.

A recent report suggested that tyrosine-protein kinase receptor AXL mediates SARS-CoV-2 entry in an ACE2-independent manner (Wang et al., 2021). It is unlikely that AXL is the alternative receptor utilized by SARS-CoV-2 in H522 cells for the following reasons. First, we found that AXL expression was lower in H522s compared with the other cell lines in our panel (**Table S1**). Second, while AXL expression enhanced lentiviruses pseudotyped with SARS-CoV-2 S (Wang et al., 2021), H522 cells were resistant to chimeric VSV-SARS-CoV-2 S and lentiviruses pseudotyped with SARS-CoV-2 S (**Fig. 2D and data not shown**). Third, though AXL potently enhances SARS-CoV-2 S pseudotyped lentivirus entry, its effect on fully infectious SARS-CoV-2 replication is lower.

Despite the possible utilization of an alternative receptor, SARS-CoV-2 entry into H522 cells requires S. However, our data suggests that S is either insufficient or that the 21 residue truncation of the cytoplasmic tail influences the ability of S to mediate infection of H522 cells, given the inability of the VSV-GFP-SARS-CoV-2-S_Δ21_ to infect H522 cells. VSV is highly sensitive to type I IFNs and numerous ISGs have been documented to block VSV replication (Espert et al., 2003; Fensterl et al., 2012; Liu et al., 2012; Muller et al., 1994; Pavlovic et al., 1990; Rihn et al., 2019; Rubinstein et al., 1981; Zurcher et al., 1992). The ability of VSV-GFP-SARS-CoV-2-S_Δ2_1 to infect H522 cells upon ACE2 expression argues against the possibility of intrinsic and IFN-induced innate immune factors targeting VSV in H522 cells. Whether the viral coat E and M proteins in SARS-CoV-2 functionally impact infection of H522 remains to be tested (Masters, 2006; Schoeman and Fielding, 2019), as it is plausible that the steric hindrance caused by blockage of S may have interfered with E/M-mediated entry.

Another important observation in our study is the inability of numerous lung and head/neck cancer cell lines to support SARS-CoV-2 replication, despite expressing ACE2/TMPRSS2. The one exception was the HCC287 cells, which express ACE2/TMPRSS2 but were less permissive to infection compared to the H522 and Vero E6 cells. One possible explanation for the general lack of permissiveness of these cell lines to SARS-CoV-2 include the alternative glycosylation or other post translations modifications of ACE2 (**Fig. 1C**). Alternatively, presence or induction of antiviral mechanisms such as type I/III IFNs and ISGs may underlie the lack of SARS-CoV-2 replication in these cells. These results collectively suggest that expression of ACE2/TMPRSS2 at endogenous levels is a poor predictor of permissiveness to SARS-CoV-2 infection.

While permissive, SARS-CoV-2 infection in H522 cells proceeded at a slower rate compared to the highly permissive Vero E6 cells. We posit that the lower susceptibility of H522 cells may be explained by the ability to mount a substantial type I IFNs response upon infection (**Fig. 5–7**) as compared to Vero E6 cells, which are unable to synthesize type I IFNs. This hypothesis is in line with previous studies demonstrating the potent inhibition of SARS-CoV-2 replication by type I IFN treatment *in vitro* (Lokugamage et al., 2020; Xie et al., 2020). Furthermore, similar to Calu-3 cells (Yin et al., 2021), we found clear involvement of MDA5 and MAVS in induction of the IFN response in H522 cells. Despite the marked activation of the type I IFN responses, virus spread in H522 cells suggests the effective antagonism of these antiviral responses, possibly through the actions of numerous viral proteins (Park and Iwasaki, 2020).

Several CoV proteins have well described functions in modulation of host gene expression post-transcriptionally (Huang et al., 2011; Kamitani et al., 2009; Kopecky-Bromberg et al., 2006; Lokugamage et al., 2012; Nakagawa et al., 2016; Narayanan et al., 2008; Xiao et al., 2008; Zhou et al., 2008). For example, SARS-CoV-2 Nsp1 is thought to block host mRNA translation through its direct binding to host ribosomes (Schubert et al., 2020; Yuan et al., 2020), which may explain the general discordance of host responses between existing RNA-seq and proteomics studies in SARS-CoV-2 infected cells (Blanco-Melo et al., 2020; Bojkova et al., 2020; Bouhaddou et al., 2020; Chu et al., 2020a; Mick et al., 2020; Stukalov et al., 2020). Similarly, we see a wide range of anti-correlated and correlated regulated genes from our transcriptomic and proteomic datasets (**Fig. 6H**), further supporting a role for SARS-CoV-2 in modulating the host cell translational response, possibly through the action of Nsp1.

Our data implicates CME in SARS-CoV-2 infection of H522 cells. Specific inhibition of a kinase directly involved in CME, AAK1, significantly reduced SARS-CoV-2 infection in H522 cells and patient-derived HBECs (**Fig. 4, S4**). Moreover, SARS-CoV-2 infection of H522 cells resulted in transient increase in phosphorylation of AP2M1, the downstream target of AAK1 kinase activity. Though inhibition of AAK1 and CME has been suggested for therapeutic treatment of Covid-19, their efficacy remains to be established. Interestingly, AAK1 inhibition preferentially blocked SARS-CoV-2 infection in H522 cells as compared to Vero E6 or Calu-3 cells (**Fig. S4**), suggesting the ACE2-independent entry mechanism in H522 cells relies on CME.

In line with published studies (Bouhaddou et al., 2020; Ochsner et al., 2020), our findings also indicate that SARS-CoV-2 may modulate DNA replication and cell cycle. However, our findings differ from these published studies in the following ways. First, instead of downmodulation of E2F targets (Ochsner et al., 2020), we find upregulation of E2F targets forming different temporal clusters (**Fig. 6E, F**). Second, in contrast to a phospho-proteomics study conducted in Vero E6 cells which only found activation of kinases involved in cell cycle regulation and DNA replication without any protein level changes (Bouhaddou et al., 2020), we find RNA and protein level upregulation of these gene sets including CCNB1,CCNE1, CHEK1, PLK1, AURKA and PKMYT1.

Notwithstanding, modulation of cell cycle upon SARS-CoV-2 is common to all of these studies, although the relevance for SARS-CoV-2 pathogenesis remains unclear.

Taken together, H522 cells provide an alternative *in vitro* model to study SARS-CoV-2 infection and host innate immune responses. The independence of virus replication from ACE2/TMPRSS2 in these cells indicates the utilization of an alternative receptor and entry pathway which may have functional relevance in understanding disease pathogenesis *in vivo*. Characterization of these mechanisms may provide unique targets for therapeutic development and vaccine design. The inevitable emergence of novel coronaviruses utilizing variable entry pathways further underscores the importance of the H522 cell line model.

## ACKNOWLEDGEMENTS

This work was supported in part by a V Foundation grant (T2014-009) to M.B. Major and D. N. Hayes, a T32 training grant (T32CA009547-34) to K.L., the Dorothy R. and Hubert C. Moog Professor of Medicine to S.L.B, a K08HL150223 grant to A.H., NIH AI059371 to S.P.J.W. We thank the Alvin J. Siteman Cancer Center at Washington University School of Medicine and Barnes-Jewish Hospital in St. Louis, MO., for the use of the Siteman Flow Cytometry, which provided single cell sorting. The Siteman Cancer Center is supported in part by an NCI Cancer Center Support Grant #P30 CA091842. We thank Dr. Ali Ellebedy for providing the 2b04 neutralizing antibody and Dr. Daved Fremont for the soluble Fc-ACE2. We additionally thank members of the Whelan and Diamond labs for reagents.

## AUTHOR CONTRIBUTIONS

M. P-C., K.L., M.B.M. and S.B.K. conceptualized the study; M. P-C., K.L., M.B.M. and S.B.K. designed the methodology. M. P-C., K.L., J.L. E., D.B., M.J.A., D.Q.L., K.D., K.T., J.E.E., C.S.M., H.R.V. and S.B.K performed the experiments. T.S., R.J., H.J., and D.G. performed all statistical and bioinformatics analysis with help from N.L. and K.T. P.W.R. and A.B. generated and provided key reagents. A.H. and S.L.B. generated and cultured primary basal epithelial cells. K.L., M.B.M. and S.B.K. wrote the manuscript with input from all the authors.

## STAR METHODS

### Resource Availability

#### Lead Contact

Further information and requests for reagents and resources should be directed to and will be fulfilled by the Lead Contact, Sebla B. Kutluay.

#### Materials Availability

All unique reagents generated in this study are available from the Lead Contact

#### Data and code availability

Raw RNA sequencing data are available on the GEO repository (GSE163547) and NCBI SRA (bioproject, PRJNA523380 and PRJNA533478) for the lung and head/neck cancer cell lines.

Raw proteomics data are available via ProteomeXchange with identifier PXD023754. Reviewer account details:

**Password:** b2aH27kS

R scripts to process data and generate figures are available on GitHub: https://github.com/GoldfarbLab/H522_paper_figures

### Experimental Model and Subject Details

#### Viral Strains

SARS-CoV-2 strain 2019-nCoV/USA-WA1/2020 was obtained from Centers for Disease Control and Prevention (a gift of Natalie Thornburg). SARS-CoV-2 was propagated in Vero CCL-81 cells (America Type Culture Collection (ATCC)-CCL-81) at an MOI of 0.01 grown in Dulbecco’s Modified Eagle’s Medium (DMEM, Sigma), supplemented with 10% Fetal Bovine Serum (FBS, VWR) and 10 mM HEPES buffer (Corning). After amplification, the virus was titered on Vero E6 cells (ATCC-CRL1586) by plaque assays and sequenced to confirm identity. E484D (100%) and R682W substitutions (10-40%) were found in our virus stocks, but we did not observe selection of additional S mutations following growth in H522 cells. All experiments involving SARS-CoV-2 were performed in a biosafety level 3 laboratory.

VSV-GFP-SARS-CoV-2-S _Δ21_ virus was kindly provided by Dr. Sean Whelan (Washington University, St. Louis) and used as previously described (Case et al., 2020). Briefly, the VSV-GFP-SARS-CoV-2-S _Δ21_ virus was propagated in the MA104 cell line (ATCC-CRL-2378.1) cultured in Medium 199 (Gibco) supplemented with 10% FBS, 1% penicillin–streptomycin, and 20 mM HEPES pH 7.7. MA104 cells were infected at an MOI of 0.01 at 37°C. After 1 hour, the media was replaced with Medium 199 supplemented with 10% FBS and 1% penicillin–streptomycin and grown at 34°C. The viral supernatant was collected 48 hpi and cell debris was cleared by centrifugation for 7.5 mins at 1000 x g. All experiments involving VSV-GFP-SARS-CoV-2-S _Δ21_ were done in a biosafety level 2 laboratory.

#### Cell Culture

All cell lines were maintained in a humidified incubator at 37°C with 5% CO_2_ unless otherwise indicated. Cell line identities were validated by short tandem repeat analysis (LabCrop, Genetica Cell Line Testing) and cultures were regularly tested for mycoplasma contamination using the MycoAlert mycoplasma detection kit (Lonza). The KYSE30 and SCC25 cell lines were a kind gift from the John Hayes Lab (UTHSC). The Vero CCL81 and Vero E6 cells were cultured in DMEM, supplemented with 10% FBS and 10 mM HEPES buffer. The A427 (kind gift from Bernard Weissman Lab (UNC)) and Detroit562 (ATCC-CCL-138) cell lines were cultured in Eagle’s Minimum Essential Medium (EMEM, Corning), supplemented with 10% FBS (Sigma), 1% penicillin–streptomycin (Corning), and 2 mmol/L □-glutamine (Gibco). The SCC25 cell line was cultured in DMEM:F12 (Corning) supplemented with 10% FBS (Sigma), 1% penicillin–streptomycin (Corning), and 400ng/ml hydrocortisone (Sigma). The H522 (ATCC-CRL-5810), H596 (ATCC HTB-178), H1299 (ATCC CRL-5803), HCC827 (ATCC CRL-2868), PC-9 (Sigma #90071810), KYSE30 (kind gift from Luke Chen (NCCU)), and OE21 (Sigma # 96062201) cell lines were cultured in RPMI-1640 (Corning) supplemented with 10% FBS (Sigma) and 1% penicillin–streptomycin (Corning). HEK293T and Calu-3 (ATCC-HTB-55) cells were cultured in DMEM (Sigma), supplemented with 10% FBS (VWR). PgsA-745 (ATCC #CRL 2242) cells were cultured in DMEM/nutrient mixture F-12 Ham, supplemented with 10% FBS (VWR).

Human airway epithelial cells were isolated from surgical excess of tracheobronchial segments of lungs donated for transplantation as previously described and were exempt from regulation by US Department of Health and Human Services regulation 45 Code of Federal Regulations Part 46 (Horani et al., 2012). Tracheobronchial cells were expanded in culture, seeded on supported membranes (Transwell; Corning, Inc.), and differentiated using ALI conditions as detailed before (Horani et al., 2018; You et al., 2002).

#### hACE2 cloning

The *Homo sapiens* angiotensin-converting enzyme 2 (ACE2), transcript variant 2 amino acid sequence (NCBI Reference Sequence: NM_021804.3) was reverse translated using the Sequence Manipulation Suite and codon optimized using Integrated DNA Technologies’ Codon Optimization Tool. This fragment was synthesized as a gene block (IDT), with 5’-TTTTCTTCCATTTCAGGTGTCGTGAGGATCC added to the 5’ end and 5’-TGAGAATTCCTCGAGGGCGGCCGCTCTAGAGTC added to the 3’ end. This product was then inserted into the pLV-EF1a-IRES-Puro vector (Addgene Plasmid #85132) that had been digested with EcoRI and BamHI using Gibson Assembly (NEB). The sequence of the resulting construct was confirmed by Sanger sequencing and propagated in Stbl3 *E. coli* cells (Life Technologies) at 30°C followed by MaxiPrep (Qiagen).

#### Lentivirus production and transduction

ACE2 expressing H522 and primary basal airway epithelial cells were generated as follows. Recombinant lentivirus was produced in HEK293T cells using a vector that expresses ACE2 driven by EF1 and a cassette to confer puromycin resistance together with psPAX2 packaging (Addgene #12260) and VSV-G envelope plasmids (Addgene #12259) as described (Horani et al., 2013). H522 and basal epithelial cells were incubated with virus-containing medium for 24 h, expanded for 3 days, then selected in puromycin (2.5 μg/mL) for 3 days.

#### SARS-CoV-2 infections, plaque assays and FACS

Prior to infection, cells were seeded at 70-80% density. Infections were done by addition of virus inoculum in cell culture media supplemented with 2% FBS and intermittent rocking for 1 h. Virus inoculum was removed, cells washed twice with 1x phosphate-buffered saline (PBS) and plated in cell culture media containing 10% FBS. Infections were monitored by plaque assays and Q-RT-PCR in cell culture supernatants. Briefly, for plaque assays, Vero E6 cells were challenged with 10-fold serial dilutions of virus-containing supernatant, incubated for 1 h at 37°C with intermittent rocking, followed by addition of 2% methylcellulose and 2X MEM containing 4% FBS. 3 days post infection cells were fixed by 4% paraformaldehyde (PFA) and stained with crystal violet solution. For plaque assays and focus forming assays (FFA) in H522 cells, 2% methylcellulose and 2X RPMI containing 10% FBS combination was used. For FACS, cells were detached then fixed with 4% PFA for 20 min at room temperature, followed by permeabilization using 0.5% Tween-20 in PBS for 10 min. Cells were blocked with 1% bovine serum albumin (BSA) and 10% FBS in 0.1% Tween-20 PBS (PBST) for 1 h prior to staining with a rabbit polyclonal anti SARS-CoV-2 nucleocapsid antibody (Sino Biological Inc. catalog # 40588-T62) diluted 1:500 and incubated overnight at 4°C. The following day, after washed cells were stained with an Alexa Fluor 488-conjugated goat anti-rabbit secondary antibody (Invitrogen) at 1:1000 dilution. FACS was performed using a BD LSR Fortessa flow cytometer and analyzed by FlowJo software. For FFA, attached cells were fixed, and stained as described for FACS, but for permeabilization 0.1% Triton-X100 was used, images were analyzed using biomolecular imager Typhoon and ImageJ software.

#### RNA extraction, qRT-PCR, and RNA-seq

Cell associated RNA was extracted by Zymo RNA-clean and concentrator-5 kit following lysis of infected cells in 1X lysis buffer (20 mM TrisHCl, 150 mM NaCl, 5 mM MgCl2, 1% Triton X-100, 1 mM DTT, 0.2 U/μL SuperaseIN RNase Inhibitor, 0.1% NP-40) and following the manufacturer’s instructions, or by Trizol extraction (Thermo Fisher Scientific). Extracted RNA was either subjected to Q-RT-PCR analysis for viral RNAs, cellular RNA, or RNA-seq. Viral RNA in cell culture supernatants was quantitated as detailed before (Shema Mugisha et al., 2020b). In brief, 5 μL of supernatant was mixed with 5 μL of 2x lysis buffer (2% Triton X-100, 50mM KCl, 100mM TrisHCl pH7.4, 40% glycerol supplemented with 400u/mL of SuperaseIN (Life Technologies)), followed by addition of 90 μL of 1X core buffer (5 mM (NH4)2SO4, 20 mM KCl and 20 mM Tris–HCl pH 8.3). 10 μL of this sample was used in a TaqMan-based Q-RT-PCR assay using TaqMan™ RNA-to-CT™ 1-Step Kit (Applied Biosystems, #4392938), alongside with RNA standards, targeting SARS-CoV-2 N gene. The primers and probe sequences are as described before (Shema Mugisha et al., 2020b). To study the interferon (IFN) response, cellular RNA was reverse transcribed with High-Capacity cDNA Reverse Transcription kit (Thermo Fisher Scientific) followed by Q-RT-PCR analysis using PowerUp SYBR Green Master Mix (Applied Biosystems). RNA levels were quantified using the ΔC_T_ method with *18S rRNA* as the reference target. ISG-specific primers are listed in Table S6.

RNA from human lung and airway cell lines were extracted using the PureLink RNA Mini Kit (Invitrogen). RT-PCR was performed on 1μg of RNA using the iScript™ gDNA Clear cDNA Synthesis Kit (Bio-Rad) and analyzed by qPCR using PowerUp SYBR Green Master Mix (Applied Biosystems) on a QuanStudio 5 machine. RNA levels were quantified using the ΔC_T_ method with *RPL13a* as the reference target. Gene specific primers are listed in Table S6.

Samples were prepared for RNA-seq using the Truseq stranded mRNA kit (Illumina) and subjected to sequencing on a Next-seq platform (1×75bp) at the Center for Genome Sciences at Washington University.

#### Immunofluorescence, RNA-ISH, and transmission electron microscopy

SARS-CoV-2 RNA and N protein were visualized in infected cells according to the published multiplex immunofluorescent cell-based detection of DNA, RNA and Protein (MICDDRP) protocol (Puray-Chavez et al., 2017). H522 cells were plated on 1.5 mm collagen-treated coverslips (GG-12-1.5-Collagen, Neuvitro) placed in 24-well plates one day prior to infection. Cells were infected with SARS-CoV-2 as above and fixed with 4% PFA at various time points post infection. Following fixation, cells were dehydrated with ethanol and stored at −20°C. Prior to probing for vRNA, cells were rehydrated, incubated in 0.1% Tween in PBS for 10 min, and mounted on slides. Probing was performed using RNAScope probes and reagents (Advanced Cell Diagnostics.) Briefly, coverslips were treated with protease solution for 15 min in a humidified HybEZ oven (Advanced Cell Diagnostics) at 40 °C. The coverslips were then washed with PBS and pre-designed anti-sense probes specific for SARS-CoV-2 positive strand S gene encoding the spike protein (RNAscope Probe-V-nCoV2019-S, cat# 848561) were applied and allowed to hybridize with the samples in a humidified HybEZ oven at 40 C for 2 hr. The probes were visualized by hybridizing with preamplifiers, amplifiers, and finally, a fluorescent label. First, pre-amplifier 1 (Amp 1-FL) was hybridized to its cognate probe for 30 min in a humidified HybEZ oven at 40 °C. Samples were then subsequently incubated with Amp 2-FL, Amp 3-FL, and Amp 4A-FL for 15 min, 30 min, and 15 min respectively. Between adding amplifiers, the coverslips were washed with a proprietary wash buffer. After probing for vRNA, samples were immunostained for the viral N protein. Coverslips were incubated in 1% bovine serum albumin (BSA) and 10% FBS in PBS containing 0.1% Tween-20 (PBST) at room temperature for 1□h. Samples were then incubated in a primary rabbit polyclonal SARS-CoV-2 nucleocapsid (N) antibody (Sino Biological Inc., Cat # 40588-T62) at 4□°C overnight. After washing in PBST, the samples were then incubated in a goat anti-rabbit fluorescent secondary antibody (Invitrogen Alexa Fluor Plus 680, Cat# A32734) at room temperature for 1 h. Nuclei were stained with DAPI diluted in PBS at room temperature for 5□min. Finally, coverslips were washed in PBST followed by PBS and then mounted on slides using Prolong Gold Antifade.

Images were taken using a Zeiss LSM 880 Airyscan confocal microscope equipped with a ×63/1.4 oil-immersion objective using the Airyscan super-resolution mode. Images were taken of the samples using either the ×63 or ×10 objective.

For ultrastructural analyses by transmission electron microscopy, samples were fixed in 2% paraformaldehyde/2.5% glutaraldehyde (Polysciences Inc., Warrington, PA) in 100 mM sodium cacodylate buffer, pH 7.2 for 1 h at room temperature. Samples were washed in sodium cacodylate buffer and postfixed in 1% osmium tetroxide (Polysciences Inc., Warrington, PA) for 1 h. Samples were then rinsed extensively in dH_2_O prior to en bloc staining with 1% aqueous uranyl acetate (Ted Pella Inc., Redding, CA) for 1 h. After several rinses in dH_2_O, samples were dehydrated in a graded series of ethanol and embedded in Eponate 12 resin (Ted Pella Inc., Redding, CA). Sections of 95 nm were cut with a Leica Ultracut UCT ultramicrotome (Leica Microsystems Inc., Bannockburn, IL), stained with uranyl acetate and lead citrate, and viewed on a JEOL 1200 EX transmission electron microscope (JEOL USA Inc., Peabody, MA) equipped with an AMT 8 megapixel digital camera and AMT Image Capture Engine V602 software (Advanced Microscopy Techniques, Woburn, MA).

#### Live cell imaging and quantification

VSV-SARS-CoV-2-S _Δ21_ viral infection rates were imaged and quantified using the Incucyte^®^ S3 Life Cell Analysis System. VeroE6, H522, and H522-ACE2 cells were labeled with Incucyte^®^ NucLight Red (Sartorius #4625) to generate stable expression of the red nuclear marker. Basal airway epithelial cells (AEC) were labeled with Incucyte^®^ NucLight Rapid Red (Sartorius #4717) at the time of infection. VSV-SARS-CoV-2-S _Δ21_ was then added to cells and immediately placed in the Incucyte^®^. Phase and fluorescent images were taken every hour to track viral infection. Percentage of GFP positive cells was calculated by dividing green object count by red object count for each well.

#### Immunoblotting

Human cell lines were grown to 70% confluence and lysed in RIPA (10% glycerol, 50mM Tris-HCl pH 7.4, 150mM NaCl, 2mM EDTA, 0.1% SDS, 1% NP40, 0.2% sodium deoxycholate) containing protease and phosphatase inhibitors (Thermo Scientific). Proteins were separated by SDS-PAGE, transferred to a nitrocellulose membrane, blocked in 5% milk, and incubated with primary antibodies overnight at 4°C. Washed membranes were incubated for 45 min at room temperature in secondary antibody solution (LI-COR IRDye 680, 800; 1:10,000 in 5% milk), imaged on an Odyssey^®^ CLx, and analyzed using Image Studio Software. Antibodies were used at the following dilutions: ACE2 (R&D Systems #AF933, 1:200), β-actin (Sigma #A5316, 1:5000), Vinculin (Santa Cruz #sc-73614, 1:2000), AAK1 (Bethyl #A302-146A, 1:1000), AP2M1 (Abcam #ab75995, 1:1000), pAP2M1-T156 (Cell Signaling #3843, 1:1000), SARS-CoV-2-N (Sino Biological #40588-T62, 1:500), pSTAT1-Y701 (Cell Signaling #9167, 1:1000), pSTAT1-S727 (Cell Signaling #8826, 1:1000), STAT1 (Cell Signaling #14994, 1:1000), MX1 (Cell Signaling #37849, 1:1000), IFIT1 (Cell Signaling #14769, 1:1000), pIKKα/β-S176/180 (Cell Signaling #2697, 1:1000), IKKα (Cell Signaling #11930, 1:1000), IKKβ (Cell Signaling #8943, 1:1000), pNFKB p65-S536 (Cell Signaling #3033, 1:1000), NFKB p65 (Cell Signaling #8242, 1:1000),

#### ACE2 CRISPR KO

H522 cells were transduced with a pLentiCRISPRv2 derived vector targeting *ACE2*(Genscript). The sgRNA (GTACTGTAGATGGTGCTCAT) targets exon 3 of *ACE2*(CCDS14169). After transduction, cells were selected in 3 μg/mL puromycin. Editing efficiency in the polyclonal population was determined using the Genomic Cleavage Detection assay (Invitrogen #A24372) and following the manufacturers protocol. The region surrounding the cleavage site was amplified using ACE2_Screen-F and ACE2_Screen-R with the sequences listed in Table S6. H522 ACE2 KO monoclonal populations were generated by limiting dilution in 96-well plates and confirmed by sanger sequencing using the above ACE2_Screen primers.

#### siRNA transfections

H522 cells were reverse-transfected by siRNAs targeting Toll-like receptor and RIG-I-like receptor pathway components. The complete sequence and catalog numbers for each siRNA are listed in Table S6. In brief, 2.5 pmoles of two siRNAs for each target was combined (total of 5 pmoles) and complexed with 0.75 μL of RNAimax transfection reagent following manufacturer’s recommendations and added in 24-well cell culture dishes. H522 cells were then seeded at 1×10^5^ cells/well. Transfected cells were infected 1-day post-transfection with SARS-CoV-2 virus, and RNA extracted 3 or 4 days later by Trizol and processed for Q-RT-PCR analysis.

#### Pharmacological effects on SARS-CoV-2 infections

In experiments where the mechanism of viral entry was probed, cells were pretreated with compounds, antibodies and ACE2-Fc as indicated in figure legends. Following viral adsorption, cells were continually kept in the presence of compounds till harvesting of the cell-associated total RNA by Trizol extraction.

#### Whole cell proteomics sample preparation

The whole cell protocol was generally followed as described before (Mertins et al., 2018). Briefly, H522 cells were grown to 80-90% confluency in 10cm cell culture dishes and then infected with SARS-CoV-2 (MOI=1 pfu/cell) for indicated time points. Cells were then lysed in urea lysis buffer containing 8 M urea, 75 mM NaCl, 50 mM Tris (pH 8.0), 1 mM EDTA, phosphatase and protease inhibitors. Samples were high speed cleared for 15 min at max speed and protein concentration was determined via BCA. 1mg of protein lysate was aliquoted, then reduced with 5mM DTT and alkylated with 15mM chloroacetamide. The sample was then diluted with 50 mM Tris-HCl (pH 8.0) to decrease the urea concentration to <2 M. Lysyl endopeptidase (Wako Chemicals, 12902541) was then added at a 1mAU to 50μg of protein and incubated at 30°C for 4 h. Trypsin (Promega, PR-V5113) was then added in an enzyme/substrate ratio of 1:49 (wt/wt) for overnight digestion at 30°C. The reaction was quenched by acidifying the mixture to a concentration of 1% formic acid. Peptides were desalted using a 200-mg tC18 SepPak cartridge (Waters Technologies, WAT054925) with a vacuum manifold (protocol followed exactly as described in (Mertins et al., 2018)) and speedvac’d dry. The sample was then resuspended in 50 mM HEPES (pH 8.5). The peptide concentration was determined by a quantitative fluorometric peptide assay kit (Pierce, PI23290). TMT labeling protocol from Zecha J, et al. was followed, except for labeling duration (Zecha et al., 2020). Briefly, 300μg of peptide was aliquoted for TMT labeling and brought to a total volume of 60μL. TMT labels (Thermo, PIA37725) were resuspended in anhydrous acetonitrile so that the concentration of each label is 20μg/μL. 15μL of label (300μg) was added to the sample and incubated at 25°C for 6 h. A test mix for labeling efficiency and label abundance was analyzed by mass spectrometry prior to mixing the samples. Samples were mixed according to the ratio determined by the test mix. The mixed sample was then desalted using a 200-mg tC18 SepPak cartridge (Waters Technologies, WAT054925) with a vacuum manifold (protocol followed exactly as described in (Mertins et al., 2018)), speedvac’d dry, and resuspended in HPLC compatible buffer. HPLC fractionation and fraction pooling was followed exactly as described in (Mertins et al., 2018). Following fractionation, samples were pooled to 25 fractions, speedvac’d dry, and resuspended in mass spec compatible buffer. 5% of each fraction was then analyzed by mass spectrometry for global whole cell proteomics.

#### Mass spectrometry data acquisition

Trypsinized peptides were separated via reverse-phase nano-HPLC using an RSLCnano Ultimate 3000 (Thermo Fisher Scientific). The mobile phase consisted of water + 0.1% formic acid as buffer A and acetonitrile + 0.1% formic acid as buffer B. Peptides were loaded onto a μPAC□□ Trapping column (PharmaFluidics) and separated on a 200 cm μPAC□□ column (PharmaFluidics) operated at 30°C using a 110 min gradient from 2% to 30% buffer B, followed by a 10 min gradient from 30% to 45% buffer B, flowing at 300 nL/min. Mass spectrometry analysis was performed on an Orbitrap Eclipse (Thermo Fisher Scientific) operated in data-dependent acquisition mode and used real-time sequencing (RTS) to trigger MS3 scans. MS1 scans were acquired in the Orbitrap at 120k resolution, with a 100% normalized automated gain control (AGC) target, auto max injection time, and a 375-1800 m/z scan range. MS2 targets were filtered for ≥50% precursor fit MS2, ≥2e4 signal intensity, charges 2-6, with a dynamic exclusion of 60 seconds, and were accumulated using a 1.2 m/z quadrupole isolation window. MS2 scans were performed in the ion trap at a turbo scan rate following collision induced dissociation (CID) at 35% collision energy. MS2 scans used a 100% normalized AGC target and auto max injection time. MS3 scans were trigged upon peptide identification using RTS. For RTS, the UniProtKB/Swiss-Prot human sequence database including isoforms (downloaded Aug. 2019) was appended with the SARS-CoV-2 proteome from UniProtKB and common contaminants from MaxQuant (Tyanova et al., 2016; UniProt, 2019). RTS parameters were set to a tryptic digestion with one missed cleavage, static Carbamidoemthyl cysteine modification (+57.0215) and TMT10 (+229.1629) on lysines and N-termini, and a variable oxidized methionine modification (+15.9949) and a maximum of 2 variable modifications per peptide. FDR filtering and protein close-out were enabled with a maximum of 5 peptides per protein and maximum search time of 50ms. The RTS scoring thresholds were set to Xcorr = 2.0, dCN = 0.1, and precursor ppm = 10 for all charges. MS3 scans were performed on the 10 most intense MS2 fragment ions identified by RTS and isolated using Synchronous Precursor Selection. MS3 scans used a normalized AGC target of 300%, auto max injection time, 50k resolution, 55% higher-energy collision dissociation collision energy, and 2 m/z wide MS2 isolation window. Acquisition was performed with a 2.5 second cycle time.

#### Whole cell proteomics raw data processing

Raw MS data files were processed by MaxQuant (version 1.6.16.0) with the same sequence database used for RTS during data acquisition. The following parameters were used: specific tryptic digestion with up to two missed cleavages, fixed carbamidomethyl modification, variable modifications for protein N-terminal acetylation, methionine oxidation, and asparagine deamidation, match between runs, and reporter ion MS3 quantification. Lot specific impurities were used for the TMT labels.

### Quantification and Statistical Analysis

Statistical parameters and details for each experiment are reported in respective figure legends. Generally, experiments were repeated with at least three biological replicates, represented by n. Each plot includes points for individual biological replicates and mean ± SEM error bars unless otherwise specified.

GraphPad Prism 9 software was used for statistical analysis. Two-way ANOVA was performed to assess significance and post hoc comparisons were employed using the Dunnett or Tukey test to correct for multiple comparisons. For statistical analysis of viral RNA (copies/cell), the data was log transformed prior to performing two-way ANOVA. The R statistical programming language was used for data processing and figure generation.

#### RNA-seq data

In order to standardize RNA-seq data from 3 different protocols in Figure 1A and Figure S1, one of which used single end 50 base reads, all reads were trimmed to 50 bp length with FASTX-Toolkit (v0.0.13) and only the reads of the first pair were considered to adjust varying read lengths and technology. Those trimmed reads were mapped to the GRCh38 genome, aided with the Gencode v35 annotation of the transcriptome with STAR (v2.7.0.f_0328). Gene expression was quantified with Salmon (v1.3.0) in alignment-based mode (Patro et al., 2017). The resulting counts were normalized to a fixed upper quartile.

Salmon v1.1.0 was used for quantification of H522 RNA-seq data (Figure 5). Salmon indexes were constructed for both hg38, Gencode v27; as well as Sars-CoV-2 based on reference genome NC_045512.2. The R tximport package was used for per gene aggregation of human transcripts based on quantification from Salmon, using Gencode v27 as well (Soneson et al., 2015). Relative expression SARS-CoV-2 RNA was expressed in terms of the log transformed ratio of total reads mapping to SARS-CoV-2 vs. hg38. 95% confidence intervals were calculated based on the assumption of normally distributed error.

Principle component analysis was performed without low-expressed genes. Starting with gene level read count quantification from Salmon and tximport as above. Genes with more than 5 counts-per-million in at least 24/48 samples were selected as an initial pre-processing step. Data normalization was performed by using the trimmed mean of M-values methods as implemented in the calcNormFactors function from the edgeR package. Normalized read counts were converted to counts-per-million and log_2_ transformed (logCPM). Principle component analysis was performed with the R function prcomp with data centering.

Differential expression analysis was performed starting with gene level read count quantification from Salmon and tximport as above. Marginally detected genes (less than 5 counts-per-million in less than 8/48 samples) were removed, as an initial pre-processing step. Data normalization was performed by using the trimmed mean of M-values methods as implemented in the calcNormFactors function from the edgeR package (Robinson et al., 2010). Normalized read counts were converted to counts-per-million and log_2_ transformed (logCPM). Differential expression analysis was subsequently performed using the Limma R-packages functions voom and eBayes (Ritchie et al., 2015). Mock infection was compared pairwise with post-infection time points, using data aggregated from the two highest MOIs (0.25 and 1.0) which appeared to have indistinguishable levels of SARS-CoV-2 gene expression. Multiple comparison correction was then performed based on the per-gene p.values from the eBayes function using the R package fdrTool. Log_2_ fold-change values (logFC) were utilized as estimated by the limma eBayes function.

Data from the two highest MOIs were used, as above, in order to cluster based on temporal gene expression changes. LogCPM values for replicate conditions were averaged. Data were then filtered for genes with both an adjusted p-value of < 0.005 and an absolute logFC > 2 at some time point in the experiment.

#### Whole-cell proteomics data

Protein abundances were computed by summing TMT reporter intensities for all spectra from a protein group’s razor and unique peptides. MS3 spectra were filtered to have >4e3 summed TMT reporter intensity and a non-missing value for the pooled bridge channel. Protein groups were filtered to have at least two unfiltered MS3 spectra in at least two of three replicate experiments. Missing values for protein abundances were imputed with the minimum protein intensity observed in the dataset. To correct for loading differences, protein abundances were normalized to have equal total abundance per TMT channel. To facilitate comparison between the three TMT-plexes, protein abundances were then divided by their corresponding pooled bridge abundance.

Differential expression was determined with the R Limma package. Moderated t-statistics were computed between the 96 mock and 96 hpi samples with default settings. Benjamini-Hochberg adjustment was used for multiple test correction. Significance filters of an adjusted p-value <0.05 and an absolute log_2_ fold-change > log_2_(1.3) were used. The lower fold-change threshold was employed in order to capture proteins whose differential expression peaked at other time points.

Protein-protein interactions for human proteins were extracted from BioGRID’s multi-validated dataset (downloaded Jan. 2021) (Oughtred et al., 2021). Interactions between human and SARS-CoV-2 proteins were extracted from BioGRID’s COVID-19 Coronavirus Project dataset (downloaded Jan. 2021) and filtered for interactions passing multi-validated criteria. Protein complexes were downloaded from CORUM (Giurgiu et al., 2019). Complexes were extracted for further analysis if they contained >2 proteins and >50% of the proteins were differentially expressed.

#### Cluster analysis

RNA-seq LogCPM values and protein abundances values were each converted to per-gene z-scores. Consensus clustering was then performed with the R ConsensusClusterPlus package (Wilkerson and Hayes, 2010). The non-defaults settings used were: reps=50, innerLinkage=“complete”, and finalLinkage=“ward.D2”. The optimal number of seven clusters was chosen by manual inspection of clustering quality for consensus matrices with k=1-12.

#### Gene set enrichment analysis

Over-representation of biological gene sets in individual temporal gene clusters for RNA-seq and proteomics data were investigated using the R clusterProfiler package and enricher function (Yu et al., 2012). Gene sets were downloaded from the MSIG data bank via the msigdbr R-project package, including “Hallmark”, “Reactome”, “GO:BP”, and “GO:CC.” (Jassal et al., 2020; Liberzon et al., 2015; Liberzon et al., 2011). Gene sets were considered significantly enriched in a cluster if adjusted p-values were < 0.05 for proteomics. “Hallmark” and “Reactome” gene sets were similarly queried in the RNA-seq data using clusterProfiler, with a cutoff for significance of adjusted p-value < 0.1.

Gene set enrichment analysis was performed on RNA-seq logCPM values (Figure 5G). Genes were ranked according to signal to noise ratio as defined by the Broad Institute GSEA software - (μ_a_ - μ_b_)/(σ_a_+σ_b_). Where μ is average the logCPM of a given gene under one experimental condition and σ the related standard deviation. Gene set enrichment analysis was then performed with default settings using the R-project fgsea package. Test gene-sets were downloaded from the MSIG data bank via the msigdbr R-project package. Only gene sets significantly associated with temporal expression clusters (using enrichr) in RNA-seq or whole cell proteomics data were subject to GSEA. Gene sets with adjusted p-values < 0.05 detected for at least one time point were considered significant.

## SUPPLEMENTAL INFORMATION

**Figure S1. Expression of *ACE2* across cell line models, related to Figure 1. A,** Unsupervised hierarchical clustering of upper quartile-normalized RNA-seq reads. Normalized RNA-seq reads were aligned to the GRCh38 and Vervet-African green monkey genomes and quantified with Salmon (v1.3.0). As indicated, RNA-seq data were generated at UNC or Washington University or obtained from the Sequence Read Archive (SRA). **B,** Read counts for *ACE2, GAPDH* and *ACTB* across the indicated cell models.

**Figure S2. Visualization of SARS-Cov-2 replication and spread in H522 cells, related to Figure 1. A**, Representative images of H522 cells infected with SARS-CoV-2 at the indicated time points and MOIs. H522 cells were fixed and stained for SARS-CoV-2 RNA (green) by RNAScope reagents and Nucleocapsid (N) protein (red) and imaged by confocal microscopy (n=2). **B**, Quantification of vRNA puncta in H522 cells infected with variable MOIs of SARS-CoV-2. Greater than 150 cells per sample from 5 different fields were counted in a blinded manner from a representative experiment.

**Figure S3. ACE2 knockout via CRISPR in H522 and Calu-3 cell lines, related to Figure 3. A**, Genomic Cleavage Detection Assay (Invitrogen) was performed following the manufacturer’s protocol on ACE2 WT or ACE2 KO CRISPR modified polyclonal cells. **B**, Sanger sequencing of genomic *ACE2* at exon 3. Unique monoclonal populations of H522 ACE2 KO’s were aligned to the human genome (‘Ref’; hg38). The red dashed lines indicate small deletions within exon 3 of ACE2.

**Figure S4. Comparative analysis of infection pathways in H522 and other permissive cells, related to Figure 4.** H522, H522-ACE2 and Vero E6 cells were pre-treated with bafilomycin A (vATPase inhibitor), SGC-AAK1-1 (clathrin-mediated endocytosis inhibitor), E64D (endosomal cathepsins inhibitor), apilimod (PIKfyve inhibitor), or camostat mesylate (TMPRSS2 inhibitor) for 1 h and then infected with SARS-CoV-2 in the presence of the inhibitors. Cell-associated SARS-CoV-2 RNA was detected by qRT-PCR 24 hpi and normalized to DMSO treated cells (n≥3). * indicates p<0.05, ** indicates p<0.01, and *** indicates p<0.001 compared to DMSO treated controls where significance was determined using two-way ANOVA and the Dunnett correction for multiple comparisons.

**Figure S5. Protein interaction networks of differentially expressed proteins in H522 cells infected with SARS-CoV-2, related to Figure 6.** Protein complexes of differentially expressed H522 and SARS-CoV-2 proteins. Complexes and functions were extracted from the CORUM database. The colors correspond to the whole cell proteomic clusters identified in ‘Fig. 6D’.

**Figure S6. siRNA knockdown efficiency for viral sensing pathways in H522 cells, related to Figure 7.** qRT-PCR for each gene targeted by siRNA in H522 cells. Knockdown efficiency was calculated compared to a non-targeting (NT) control. H522 cells were infected with SARS-CoV-2 24 hpi and RNA was collected 24, 96, and 120 hpi. TLR8 mRNA was not detected across the three time points.

**Table S1. Cell line RNA-seq, related to Figure 1.**

**Table S2. Differentially expressed genes from RNA-seq in H522 cells infected with SARS-CoV-2, related to Figure 5.**

**Table S3. Gene set enrichment analysis from RNA-seq in H522 cells infected with SARS-CoV-2, related to Figure 5.**

**Table S4. Protein expression changes from whole cell proteomics in H522 cells infected with SARS-CoV-2, related to Figure 6.**

**Table S5. Gene set enrichment analysis from whole cell proteomics in H522 cells infected with SARS-CoV-2, related to Figure 6.**

**Table S6. Oligo sequences, related to STAR methods**

## REFERENCES

Agajanian, M.J., Walker, M.P., Axtman, A.D., Ruela-de-Sousa, R.R., Serafin, D.S., Rabinowitz, A.D., Graham, D.M., Ryan, M.B., Tamir, T., Nakamichi, Y., et al. (2019). WNT Activates the AAK1 Kinase to Promote Clathrin-Mediated Endocytosis of LRP6 and Establish a Negative Feedback Loop. Cell Rep 26, 79–93 e78.

Aguiar, J.A., Tremblay, B.J., Mansfield, M.J., Woody, O., Lobb, B., Banerjee, A., Chandiramohan, A., Tiessen, N., Cao, Q., Dvorkin-Gheva, A., et al. (2020). Gene expression and in situ protein profiling of candidate SARS-CoV-2 receptors in human airway epithelial cells and lung tissue. Eur Respir J 56.

Alsoussi, W.B., Turner, J.S., Case, J.B., Zhao, H., Schmitz, A.J., Zhou, J.Q., Chen, R.E., Lei, T., Rizk, A.A., McIntire, K.M., et al. (2020). A Potently Neutralizing Antibody Protects Mice against SARS-CoV-2 Infection. J Immunol 205, 915–922.

Blanco-Melo, D., Nilsson-Payant, B.E., Liu, W.C., Uhl, S., Hoagland, D., Moller, R., Jordan, T.X., Oishi, K., Panis, M., Sachs, D., et al. (2020). Imbalanced Host Response to SARS-CoV-2 Drives Development of COVID-19. Cell 181, 1036–1045 e1039.

Bojkova, D., Klann, K., Koch, B., Widera, M., Krause, D., Ciesek, S., Cinatl, J., and Munch, C. (2020). Proteomics of SARS-CoV-2-infected host cells reveals therapy targets. Nature 583, 469–472.

Bouhaddou, M., Memon, D., Meyer, B., White, K.M., Rezelj, V.V., Correa Marrero, M., Polacco, B.J., Melnyk, J.E., Ulferts, S., Kaake, R.M., et al. (2020). The Global Phosphorylation Landscape of SARS-CoV-2 Infection. Cell 182, 685–712 e619.

Cagno, V. (2020). SARS-CoV-2 cellular tropism. Lancet Microbe 1, e2–e3.

Cantuti-Castelvetri, L., Ojha, R., Pedro, L.D., Djannatian, M., Franz, J., Kuivanen, S., van der Meer, F., Kallio, K., Kaya, T., Anastasina, M., et al. (2020). Neuropilin-1 facilitates SARS-CoV-2 cell entry and infectivity. Science 370, 856–860.

Case, J.B., Rothlauf, P.W., Chen, R.E., Liu, Z., Zhao, H., Kim, A.S., Bloyet, L.M., Zeng, Q., Tahan, S., Droit, L., et al. (2020). Neutralizing Antibody and Soluble ACE2 Inhibition of a Replication-Competent VSV-SARS-CoV-2 and a Clinical Isolate of SARS-CoV-2. Cell Host Microbe 28, 475–485 e475.

Chen, G., Wu, D., Guo, W., Cao, Y., Huang, D., Wang, H., Wang, T., Zhang, X., Chen, H., Yu, H., et al. (2020). Clinical and immunological features of severe and moderate coronavirus disease 2019. J Clin Invest 130, 2620–2629.

Chu, H., Chan, J.F., Wang, Y., Yuen, T.T., Chai, Y., Hou, Y., Shuai, H., Yang, D., Hu, B., Huang, X., et al. (2020a). Comparative replication and immune activation profiles of SARS-CoV-2 and SARS-CoV in human lungs: an ex vivo study with implications for the pathogenesis of COVID-19. Clin Infect Dis.

Chu, H., Chan, J.F.-W., Yuen, T.T.-T., Shuai, H., Yuan, S., Wang, Y., Hu, B., Yip, C.C.-Y., Tsang, J.O.-L., Huang, X., et al. (2020b). Comparative tropism, replication kinetics, and cell damage profiling of SARS-CoV-2 and SARS-CoV with implications for clinical manifestations, transmissibility, and laboratory studies of COVID-19: an observational study. Lancet Microbe 1, e14–e23.

Clausen, T.M., Sandoval, D.R., Spliid, C.B., Pihl, J., Perrett, H.R., Painter, C.D., Narayanan, A., Majowicz, S.A., Kwong, E.M., McVicar, R.N., et al. (2020). SARS-CoV-2 Infection Depends on Cellular Heparan Sulfate and ACE2. Cell 183, 1043–1057 e1015.

Conner, S.D., and Schmid, S.L. (2002). Identification of an adaptor-associated kinase, AAK1, as a regulator of clathrin-mediated endocytosis. J Cell Biol 156, 921–929.

Conner, S.D., and Schmid, S.L. (2003). Differential requirements for AP-2 in clathrin-mediated endocytosis. J Cell Biol 162, 773–779.

Daly, J.L., Simonetti, B., Klein, K., Chen, K.E., Williamson, M.K., Anton-Plagaro, C., Shoemark, D.K., Simon-Gracia, L., Bauer, M., Hollandi, R., et al. (2020). Neuropilin-1 is a host factor for SARS-CoV-2 infection. Science 370, 861–865.

Desmyter, J., Melnick, J.L., and Rawls, W.E. (1968). Defectiveness of interferon production and of rubella virus interference in a line of African green monkey kidney cells (Vero). J Virol 2, 955–961.

Diaz, M.O., Ziemin, S., Le Beau, M.M., Pitha, P., Smith, S.D., Chilcote, R.R., and Rowley, J.D. (1988). Homozygous deletion of the alpha- and beta 1-interferon genes in human leukemia and derived cell lines. Proc Natl Acad Sci U S A 85, 5259–5263.

Espert, L., Degols, G., Gongora, C., Blondel, D., Williams, B.R., Silverman, R.H., and Mechti, N. (2003). ISG20, a new interferon-induced RNase specific for single-stranded RNA, defines an alternative antiviral pathway against RNA genomic viruses. J Biol Chem 278, 16151–16158.

Fensterl, V., Wetzel, J.L., Ramachandran, S., Ogino, T., Stohlman, S.A., Bergmann, C.C., Diamond, M.S., Virgin, H.W., and Sen, G.C. (2012). Interferon-induced Ifit2/ISG54 protects mice from lethal VSV neuropathogenesis. PLoS Pathog 8, e1002712.

Giurgiu, M., Reinhard, J., Brauner, B., Dunger-Kaltenbach, I., Fobo, G., Frishman, G., Montrone, C., and Ruepp, A. (2019). CORUM: the comprehensive resource of mammalian protein complexes-2019. Nucleic Acids Res 47, D559–D563.

Hamming, I., Timens, W., Bulthuis, M.L., Lely, A.T., Navis, G., and van Goor, H. (2004). Tissue distribution of ACE2 protein, the functional receptor for SARS coronavirus. A first step in understanding SARS pathogenesis. J Pathol 203, 631–637.

Hikmet, F., Mear, L., Edvinsson, A., Micke, P., Uhlen, M., and Lindskog, C. (2020). The protein expression profile of ACE2 in human tissues. Mol Syst Biol 16, e9610.

Hoffmann, M., Kleine-Weber, H., Schroeder, S., Kruger, N., Herrler, T., Erichsen, S., Schiergens, T.S., Herrler, G., Wu, N.H., Nitsche, A., et al. (2020). SARS-CoV-2 Cell Entry Depends on ACE2 and TMPRSS2 and Is Blocked by a Clinically Proven Protease Inhibitor. Cell 181, 271–280 e278.

Horani, A., Druley, T.E., Zariwala, M.A., Patel, A.C., Levinson, B.T., Van Arendonk, L.G., Thornton, K.C., Giacalone, J.C., Albee, A.J., Wilson, K.S., et al. (2012). Whole-exome capture and sequencing identifies HEATR2 mutation as a cause of primary ciliary dyskinesia. Am J Hum Genet 91, 685–693.

Horani, A., Nath, A., Wasserman, M.G., Huang, T., and Brody, S.L. (2013). Rho-associated protein kinase inhibition enhances airway epithelial Basal-cell proliferation and lentivirus transduction. Am J Respir Cell Mol Biol 49, 341–347.

Horani, A., Ustione, A., Huang, T., Firth, A.L., Pan, J., Gunsten, S.P., Haspel, J.A., Piston, D.W., and Brody, S.L. (2018). Establishment of the early cilia preassembly protein complex during motile ciliogenesis. Proc Natl Acad Sci U S A 115, E1221–E1228.

Hou, Y.J., Okuda, K., Edwards, C.E., Martinez, D.R., Asakura, T., Dinnon, K.H., 3rd, Kato, T., Lee, R.E., Yount, B.L., Mascenik, T.M., et al. (2020). SARS-CoV-2 Reverse Genetics Reveals a Variable Infection Gradient in the Respiratory Tract. Cell 182, 429–446 e414.

Huang, C., Lokugamage, K.G., Rozovics, J.M., Narayanan, K., Semler, B.L., and Makino, S. (2011). SARS coronavirus nsp1 protein induces template-dependent endonucleolytic cleavage of mRNAs: viral mRNAs are resistant to nsp1-induced RNA cleavage. PLoS Pathog 7, e1002433.

Huang, C., Wang, Y., Li, X., Ren, L., Zhao, J., Hu, Y., Zhang, L., Fan, G., Xu, J., Gu, X., et al. (2020). Clinical features of patients infected with 2019 novel coronavirus in Wuhan, China. Lancet 395, 497–506.

Huang, X., Dong, W., Milewska, A., Golda, A., Qi, Y., Zhu, Q.K., Marasco, W.A., Baric, R.S., Sims, A.C., Pyrc, K., et al. (2015). Human Coronavirus HKU1 Spike Protein Uses O-Acetylated Sialic Acid as an Attachment Receptor Determinant and Employs Hemagglutinin-Esterase Protein as a Receptor-Destroying Enzyme. J Virol 89, 7202–7213.

Hulswit, R.J.G., Lang, Y., Bakkers, M.J.G., Li, W., Li, Z., Schouten, A., Ophorst, B., van Kuppeveld, F.J.M., Boons, G.J., Bosch, B.J., et al. (2019). Human coronaviruses OC43 and HKU1 bind to 9-O-acetylated sialic acids via a conserved receptor-binding site in spike protein domain A. Proc Natl Acad Sci U S A 116, 2681–2690.

Jassal, B., Matthews, L., Viteri, G., Gong, C., Lorente, P., Fabregat, A., Sidiropoulos, K., Cook, J., Gillespie, M., Haw, R., et al. (2020). The reactome pathway knowledgebase. Nucleic Acids Res 48, D498–D503.

Kamitani, W., Huang, C., Narayanan, K., Lokugamage, K.G., and Makino, S. (2009). A two-pronged strategy to suppress host protein synthesis by SARS coronavirus Nsp1 protein. Nat Struct Mol Biol 16, 1134–1140.

Kang, Y.L., Chou, Y.Y., Rothlauf, P.W., Liu, Z., Soh, T.K., Cureton, D., Case, J.B., Chen, R.E., Diamond, M.S., Whelan, S.P.J., et al. (2020). Inhibition of PIKfyve kinase prevents infection by Zaire ebolavirus and SARS-CoV-2. Proc Natl Acad Sci U S A 117, 20803–20813.

Kim, J.M., Chung, Y.S., Jo, H.J., Lee, N.J., Kim, M.S., Woo, S.H., Park, S., Kim, J.W., Kim, H.M., and Han, M.G. (2020). Identification of Coronavirus Isolated from a Patient in Korea with COVID-19. Osong Public Health Res Perspect 11, 3–7.

Kopecky-Bromberg, S.A., Martinez-Sobrido, L., and Palese, P. (2006). 7a protein of severe acute respiratory syndrome coronavirus inhibits cellular protein synthesis and activates p38 mitogen-activated protein kinase. J Virol 80, 785–793.

Lazear, H.M., Schoggins, J.W., and Diamond, M.S. (2019). Shared and Distinct Functions of Type I and Type III Interferons. Immunity 50, 907–923.

Letko, M., Marzi, A., and Munster, V. (2020). Functional assessment of cell entry and receptor usage for SARS-CoV-2 and other lineage B betacoronaviruses. Nat Microbiol 5, 562–569.

Li, W., Hulswit, R.J.G., Widjaja, I., Raj, V.S., McBride, R., Peng, W., Widagdo, W., Tortorici, M.A., van Dieren, B., Lang, Y., et al. (2017). Identification of sialic acid-binding function for the Middle East respiratory syndrome coronavirus spike glycoprotein. Proc Natl Acad Sci U S A 114, E8508–E8517.

Liberzon, A., Birger, C., Thorvaldsdottir, H., Ghandi, M., Mesirov, J.P., and Tamayo, P. (2015). The Molecular Signatures Database (MSigDB) hallmark gene set collection. Cell Syst 1, 417–425.

Liberzon, A., Subramanian, A., Pinchback, R., Thorvaldsdottir, H., Tamayo, P., and Mesirov, J.P. (2011). Molecular signatures database (MSigDB) 3.0. Bioinformatics 27, 1739–1740.

Liu, C., Tang, J., Ma, Y., Liang, X., Yang, Y., Peng, G., Qi, Q., Jiang, S., Li, J., Du, L., et al. (2015). Receptor usage and cell entry of porcine epidemic diarrhea coronavirus. J Virol 89, 6121–6125.

Liu, S.Y., Sanchez, D.J., Aliyari, R., Lu, S., and Cheng, G. (2012). Systematic identification of type I and type II interferon-induced antiviral factors. Proc Natl Acad Sci U S A 109, 4239–4244.

Lokugamage, K.G., Hage, A., de Vries, M., Valero-Jimenez, A.M., Schindewolf, C., Dittmann, M., Rajsbaum, R., and Menachery, V.D. (2020). Type I Interferon Susceptibility Distinguishes SARS-CoV-2 from SARS-CoV. J Virol 94.

Lokugamage, K.G., Narayanan, K., Huang, C., and Makino, S. (2012). Severe acute respiratory syndrome coronavirus protein nsp1 is a novel eukaryotic translation inhibitor that represses multiple steps of translation initiation. J Virol 86, 13598–13608.

Long, J.S., Mistry, B., Haslam, S.M., and Barclay, W.S. (2019). Host and viral determinants of influenza A virus species specificity. Nat Rev Microbiol 17, 67–81.

Malim, M.H., and Bieniasz, P.D. (2012). HIV Restriction Factors and Mechanisms of Evasion. Cold Spring Harb Perspect Med 2, a006940.

Masters, P.S. (2006). The molecular biology of coronaviruses. Adv Virus Res 66, 193–292.

Mertins, P., Tang, L.C., Krug, K., Clark, D.J., Gritsenko, M.A., Chen, L., Clauser, K.R., Clauss, T.R., Shah, P., Gillette, M.A., et al. (2018). Reproducible workflow for multiplexed deep-scale proteome and phosphoproteome analysis of tumor tissues by liquid chromatography-mass spectrometry. Nat Protoc 13, 1632–1661.

Mick, E., Kamm, J., Pisco, A.O., Ratnasiri, K., Babik, J.M., Castaneda, G., DeRisi, J.L., Detweiler, A.M., Hao, S.L., Kangelaris, K.N., et al. (2020). Upper airway gene expression reveals suppressed immune responses to SARS-CoV-2 compared with other respiratory viruses. Nat Commun 11, 5854.

Muller, U., Steinhoff, U., Reis, L.F., Hemmi, S., Pavlovic, J., Zinkernagel, R.M., and Aguet, M. (1994). Functional role of type I and type II interferons in antiviral defense. Science 264, 1918–1921.

Nakagawa, K., Lokugamage, K.G., and Makino, S. (2016). Viral and Cellular mRNA Translation in Coronavirus-Infected Cells. Adv Virus Res 96, 165–192.

Narayanan, K., Huang, C., Lokugamage, K., Kamitani, W., Ikegami, T., Tseng, C.T., and Makino, S. (2008). Severe acute respiratory syndrome coronavirus nsp1 suppresses host gene expression, including that of type I interferon, in infected cells. J Virol 82, 4471–4479.

Ochsner, S.A., Pillich, R.T., and McKenna, N.J. (2020). Consensus transcriptional regulatory networks of coronavirus-infected human cells. Sci Data 7, 314.

Osada, N., Kohara, A., Yamaji, T., Hirayama, N., Kasai, F., Sekizuka, T., Kuroda, M., and Hanada, K. (2014). The genome landscape of the african green monkey kidney-derived vero cell line. DNA Res 21, 673–683.

Ou, X., Liu, Y., Lei, X., Li, P., Mi, D., Ren, L., Guo, L., Guo, R., Chen, T., Hu, J., et al. (2020). Characterization of spike glycoprotein of SARS-CoV-2 on virus entry and its immune cross-reactivity with SARS-CoV. Nat Commun 11, 1620.

Oughtred, R., Rust, J., Chang, C., Breitkreutz, B.J., Stark, C., Willems, A., Boucher, L., Leung, G., Kolas, N., Zhang, F., et al. (2021). The BioGRID database: A comprehensive biomedical resource of curated protein, genetic, and chemical interactions. Protein Sci 30, 187–200.

Park, A., and Iwasaki, A. (2020). Type I and Type III Interferons - Induction, Signaling, Evasion, and Application to Combat COVID-19. Cell Host Microbe 27, 870–878.

Park, Y.J., Walls, A.C., Wang, Z., Sauer, M.M., Li, W., Tortorici, M.A., Bosch, B.J., DiMaio, F., and Veesler, D. (2019). Structures of MERS-CoV spike glycoprotein in complex with sialoside attachment receptors. Nat Struct Mol Biol 26, 1151–1157.

Patro, R., Duggal, G., Love, M.I., Irizarry, R.A., and Kingsford, C. (2017). Salmon provides fast and bias-aware quantification of transcript expression. Nat Methods 14, 417–419.

Pavlovic, J., Zurcher, T., Haller, O., and Staeheli, P. (1990). Resistance to influenza virus and vesicular stomatitis virus conferred by expression of human MxA protein. J Virol 64, 3370–3375.

Puray-Chavez, M., Tedbury, P.R., Huber, A.D., Ukah, O.B., Yapo, V., Liu, D., Ji, J., Wolf, J.J., Engelman, A.N., and Sarafianos, S.G. (2017). Multiplex single-cell visualization of nucleic acids and protein during HIV infection. Nat Commun 8, 1882.

Qi, F., Qian, S., Zhang, S., and Zhang, Z. (2020). Single cell RNA sequencing of 13 human tissues identify cell types and receptors of human coronaviruses. Biochem Biophys Res Commun 526, 135–140.

Richardson, P., Griffin, I., Tucker, C., Smith, D., Oechsle, O., Phelan, A., Rawling, M., Savory, E., and Stebbing, J. (2020). Baricitinib as potential treatment for 2019-nCoV acute respiratory disease. Lancet 395, e30–e31.

Rihn, S.J., Aziz, M.A., Stewart, D.G., Hughes, J., Turnbull, M.L., Varela, M., Sugrue, E., Herd, C.S., Stanifer, M., Sinkins, S.P., et al. (2019). TRIM69 Inhibits Vesicular Stomatitis Indiana Virus. J Virol 93.

Ritchie, M.E., Phipson, B., Wu, D., Hu, Y., Law, C.W., Shi, W., and Smyth, G.K. (2015). limma powers differential expression analyses for RNA-sequencing and microarray studies. Nucleic Acids Res 43, e47.

Robinson, M.D., McCarthy, D.J., and Smyth, G.K. (2010). edgeR: a Bioconductor package for differential expression analysis of digital gene expression data. Bioinformatics 26, 139–140.

Rothenburg, S., and Brennan, G. (2020). Species-Specific Host-Virus Interactions: Implications for Viral Host Range and Virulence. Trends Microbiol 28, 46–56.

Rubinstein, S., Familletti, P.C., and Pestka, S. (1981). Convenient assay for interferons. J Virol 37, 755–758.

Schaefer, I.M., Padera, R.F., Solomon, I.H., Kanjilal, S., Hammer, M.M., Hornick, J.L., and Sholl, L.M. (2020). In situ detection of SARS-CoV-2 in lungs and airways of patients with COVID-19. Mod Pathol 33, 2104–2114.

Schoeman, D., and Fielding, B.C. (2019). Coronavirus envelope protein: current knowledge. Virol J 16, 69.

Schoggins, J.W. (2018). Recent advances in antiviral interferon-stimulated gene biology. F1000Res 7, 309.

Schubert, K., Karousis, E.D., Jomaa, A., Scaiola, A., Echeverria, B., Gurzeler, L.A., Leibundgut, M., Thiel, V., Muhlemann, O., and Ban, N. (2020). SARS-CoV-2 Nsp1 binds the ribosomal mRNA channel to inhibit translation. Nat Struct Mol Biol 27, 959–966.

Schultze, B., Krempl, C., Ballesteros, M.L., Shaw, L., Schauer, R., Enjuanes, L., and Herrler, G. (1996). Transmissible gastroenteritis coronavirus, but not the related porcine respiratory coronavirus, has a sialic acid (N-glycolylneuraminic acid) binding activity. J Virol 70, 5634–5637.

Shajahan, A., Archer-Hartmann, S., Supekar, N.T., Gleinich, A.S., Heiss, C., and Azadi, P. (2020). Comprehensive characterization of N- and O- glycosylation of SARS-CoV-2 human receptor angiotensin converting enzyme 2. Glycobiology.

Shang, J., Wan, Y., Luo, C., Ye, G., Geng, Q., Auerbach, A., and Li, F. (2020). Cell entry mechanisms of SARS-CoV-2. Proc Natl Acad Sci U S A 117, 11727–11734.

Shema Mugisha, C., Vuong, H.R., Puray-Chavez, M., Bailey, A.L., Fox, J.M., Chen, R.E., Wessel, A.W., Scott, J.M., Harastani, H.H., Boon, A.C.M., et al. (2020a). A Simplified Quantitative Real-Time PCR Assay for Monitoring SARS-CoV-2 Growth in Cell Culture. mSphere 5.

Shema Mugisha, C., Vuong, H.R., Puray-Chavez, M., and Kutluay, S.B. (2020b). A facile Q-RT-PCR assay for monitoring SARS-CoV-2 growth in cell culture. bioRxiv.

Soneson, C., Love, M.I., and Robinson, M.D. (2015). Differential analyses for RNA-seq: transcript-level estimates improve gene-level inferences. F1000Res 4, 1521.

Stukalov, A., Girault, V., Grass, V., Bergant, V., Karayel, O., Urban, C., Haas, D.A., Huang, Y., Oubraham, L., Wang, A., et al. (2020). Multi-level proteomics reveals host-perturbation strategies of SARS-CoV-2 and SARS-CoV. bioRxiv, 2020.2006.2017.156455.

Sungnak, W., Huang, N., Becavin, C., Berg, M., Queen, R., Litvinukova, M., Talavera-Lopez, C., Maatz, H., Reichart, D., Sampaziotis, F., et al. (2020). SARS-CoV-2 entry factors are highly expressed in nasal epithelial cells together with innate immune genes. Nat Med 26, 681–687.

To, K.F., and Lo, A.W. (2004). Exploring the pathogenesis of severe acute respiratory syndrome (SARS): the tissue distribution of the coronavirus (SARS-CoV) and its putative receptor, angiotensin-converting enzyme 2 (ACE2). J Pathol 203, 740–743.

Tortorici, M.A., Walls, A.C., Lang, Y., Wang, C., Li, Z., Koerhuis, D., Boons, G.J., Bosch, B.J., Rey, F.A., de Groot, R.J., et al. (2019). Structural basis for human coronavirus attachment to sialic acid receptors. Nat Struct Mol Biol 26, 481–489.

Tyanova, S., Temu, T., and Cox, J. (2016). The MaxQuant computational platform for mass spectrometry-based shotgun proteomics. Nat Protoc 11, 2301–2319.

UniProt, C. (2019). UniProt: a worldwide hub of protein knowledge. Nucleic Acids Res 47, D506–D515.

Varga, Z., Flammer, A.J., Steiger, P., Haberecker, M., Andermatt, R., Zinkernagel, A.S., Mehra, M.R., Schuepbach, R.A., Ruschitzka, F., and Moch, H. (2020). Endothelial cell infection and endotheliitis in COVID-19. Lancet 395, 1417–1418.

Vlasak, R., Luytjes, W., Spaan, W., and Palese, P. (1988). Human and bovine coronaviruses recognize sialic acid-containing receptors similar to those of influenza C viruses. Proc Natl Acad Sci U S A 85, 4526–4529.

Walls, A.C., Park, Y.J., Tortorici, M.A., Wall, A., McGuire, A.T., and Veesler, D. (2020). Structure, Function, and Antigenicity of the SARS-CoV-2 Spike Glycoprotein. Cell 181, 281–292 e286.

Wang, S., Qiu, Z., Hou, Y., Deng, X., Xu, W., Zheng, T., Wu, P., Xie, S., Bian, W., Zhang, C., et al. (2021). AXL is a candidate receptor for SARS-CoV-2 that promotes infection of pulmonary and bronchial epithelial cells. Cell Res.

Wickramasinghe, I.N., de Vries, R.P., Grone, A., de Haan, C.A., and Verheije, M.H. (2011). Binding of avian coronavirus spike proteins to host factors reflects virus tropism and pathogenicity. J Virol 85, 8903–8912.

Wilkerson, M.D., and Hayes, D.N. (2010). ConsensusClusterPlus: a class discovery tool with confidence assessments and item tracking. Bioinformatics 26, 1572–1573.

Wrapp, D., Wang, N., Corbett, K.S., Goldsmith, J.A., Hsieh, C.L., Abiona, O., Graham, B.S., and McLellan, J.S. (2020). Cryo-EM structure of the 2019-nCoV spike in the prefusion conformation. Science 367, 1260–1263.

Wu, F., Zhao, S., Yu, B., Chen, Y.M., Wang, W., Song, Z.G., Hu, Y., Tao, Z.W., Tian, J.H., Pei, Y.Y., et al. (2020). A new coronavirus associated with human respiratory disease in China. Nature 579, 265–269.

Wu, Z., and McGoogan, J.M. (2020). Characteristics of and Important Lessons From the Coronavirus Disease 2019 (COVID-19) Outbreak in China: Summary of a Report of 72314 Cases From the Chinese Center for Disease Control and Prevention. JAMA.

Xia, H., Cao, Z., Xie, X., Zhang, X., Chen, J.Y., Wang, H., Menachery, V.D., Rajsbaum, R., and Shi, P.Y. (2020). Evasion of Type I Interferon by SARS-CoV-2. Cell Rep 33, 108234.

Xiao, H., Xu, L.H., Yamada, Y., and Liu, D.X. (2008). Coronavirus spike protein inhibits host cell translation by interaction with eIF3f. PLoS One 3, e1494.

Xie, X., Muruato, A., Lokugamage, K.G., Narayanan, K., Zhang, X., Zou, J., Liu, J., Schindewolf, C., Bopp, N.E., Aguilar, P.V., et al. (2020). An Infectious cDNA Clone of SARS-CoV-2. Cell Host Microbe 27, 841–848 e843.

Yang, Q., Hughes, T.A., Kelkar, A., Yu, X., Cheng, K., Park, S., Huang, W.C., Lovell, J.F., and Neelamegham, S. (2020). Inhibition of SARS-CoV-2 viral entry upon blocking N- and O-glycan elaboration. Elife 9.

Yin, X., Riva, L., Pu, Y., Martin-Sancho, L., Kanamune, J., Yamamoto, Y., Sakai, K., Gotoh, S., Miorin, L., De Jesus, P.D., et al. (2021). MDA5 Governs the Innate Immune Response to SARS-CoV-2 in Lung Epithelial Cells. Cell Rep 34, 108628.

You, Y., Richer, E.J., Huang, T., and Brody, S.L. (2002). Growth and differentiation of mouse tracheal epithelial cells: selection of a proliferative population. Am J Physiol Lung Cell Mol Physiol 283, L1315–1321.

Yu, G., Wang, L.G., Han, Y., and He, Q.Y. (2012). clusterProfiler: an R package for comparing biological themes among gene clusters. OMICS 16, 284–287.

Yuan, S., Peng, L., Park, J.J., Hu, Y., Devarkar, S.C., Dong, M.B., Shen, Q., Wu, S., Chen, S., Lomakin, I.B., et al. (2020). Nonstructural Protein 1 of SARS-CoV-2 Is a Potent Pathogenicity Factor Redirecting Host Protein Synthesis Machinery toward Viral RNA. Mol Cell.

Zecha, J., Lee, C.Y., Bayer, F.P., Meng, C., Grass, V., Zerweck, J., Schnatbaum, K., Michler, T., Pichlmair, A., Ludwig, C., et al. (2020). Data, Reagents, Assays and Merits of Proteomics for SARS-CoV-2 Research and Testing. Mol Cell Proteomics 19, 1503–1522.

Zhao, Y., Zhao, Z., Wang, Y., Zhou, Y., Ma, Y., and Zuo, W. (2020). Single-cell RNA Expression Profiling of ACE2, The Receptor of SARS-CoV-2. Am J Respir Crit Care Med.

Zhou, B., Liu, J., Wang, Q., Liu, X., Li, X., Li, P., Ma, Q., and Cao, C. (2008). The nucleocapsid protein of severe acute respiratory syndrome coronavirus inhibits cell cytokinesis and proliferation by interacting with translation elongation factor 1alpha. J Virol 82, 6962–6971.

Zhou, P., Yang, X.L., Wang, X.G., Hu, B., Zhang, L., Zhang, W., Si, H.R., Zhu, Y., Li, B., Huang, C.L., et al. (2020). A pneumonia outbreak associated with a new coronavirus of probable bat origin. Nature 579, 270–273.

Zou, X., Chen, K., Zou, J., Han, P., Hao, J., and Han, Z. (2020). Single-cell RNA-seq data analysis on the receptor ACE2 expression reveals the potential risk of different human organs vulnerable to 2019-nCoV infection. Front Med 14, 185–192.

Zurcher, T., Pavlovic, J., and Staeheli, P. (1992). Mouse Mx2 protein inhibits vesicular stomatitis virus but not influenza virus. Virology 187, 796–800.

