## Supplementary figures and images for "Systematic analysis of SARS-CoV-2 infection of an ACE2-negative human airway cell"

### Supplemental Figures

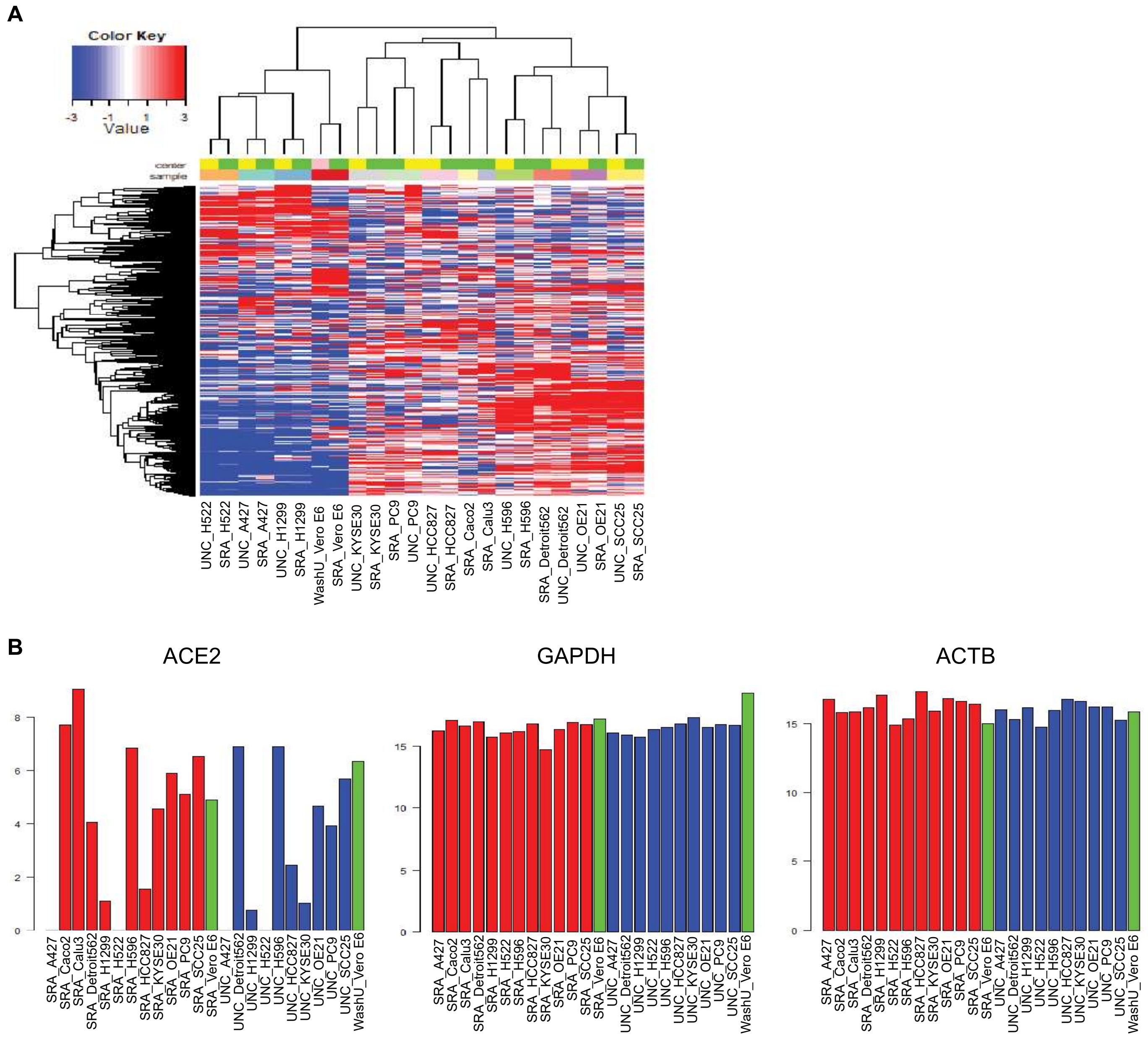

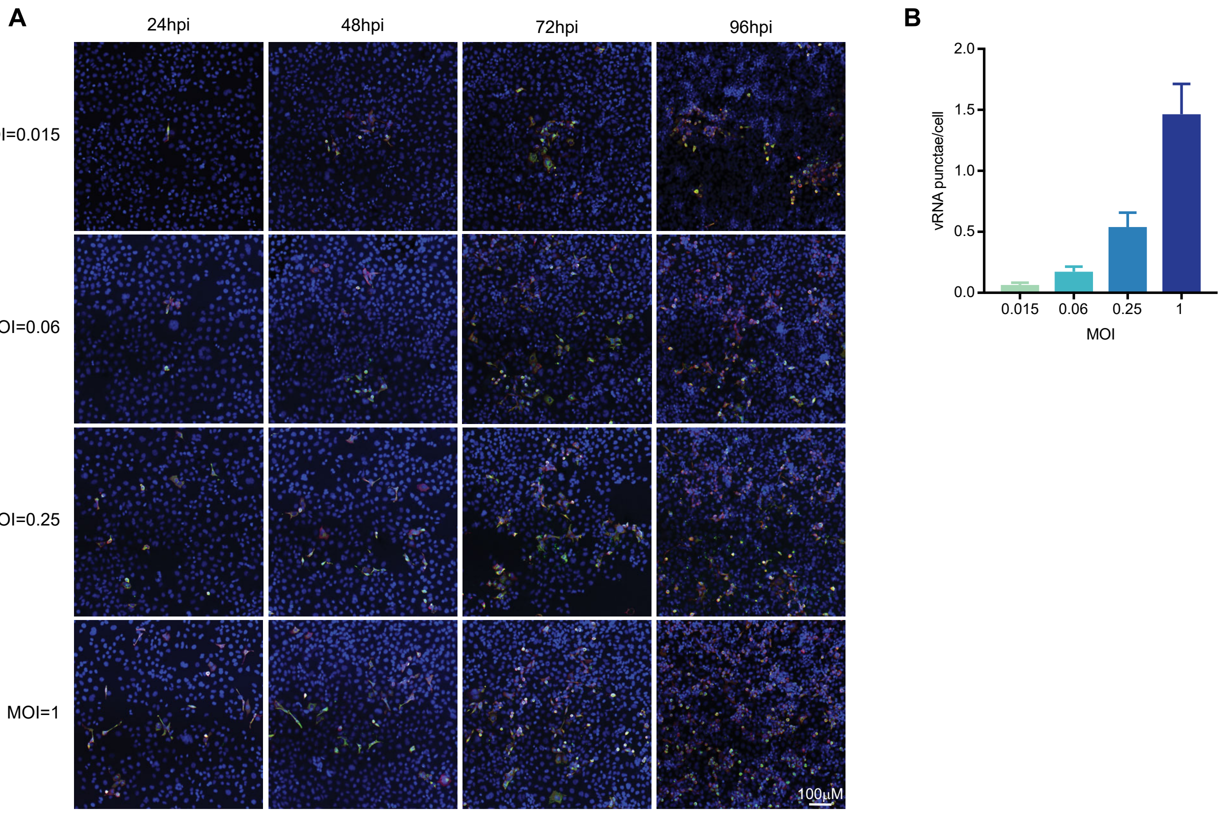

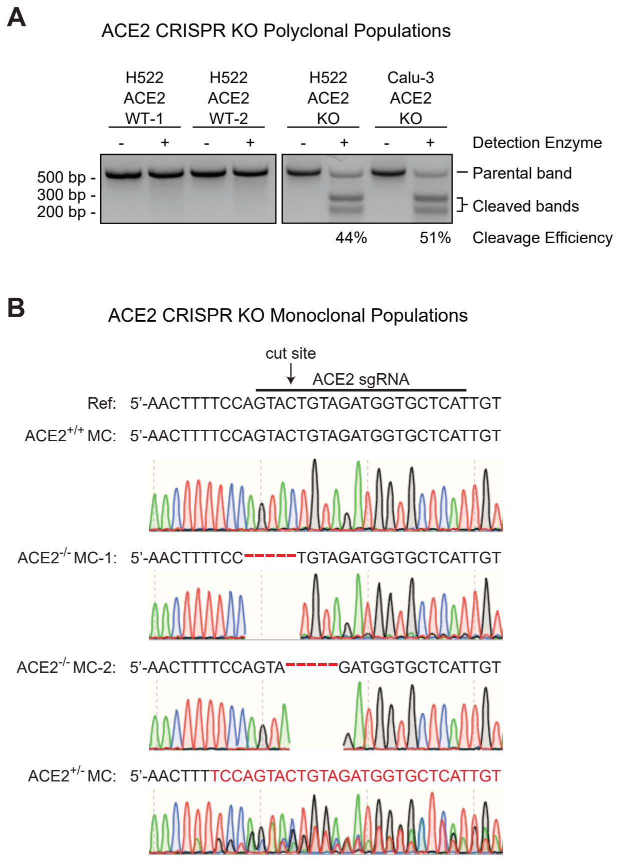

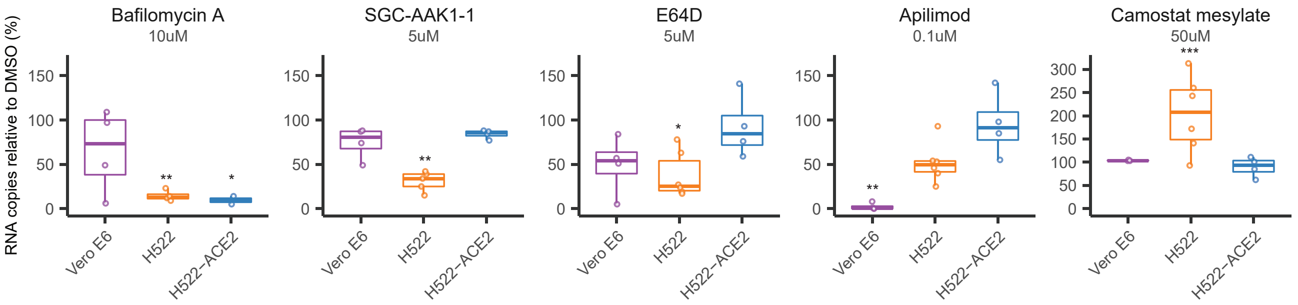

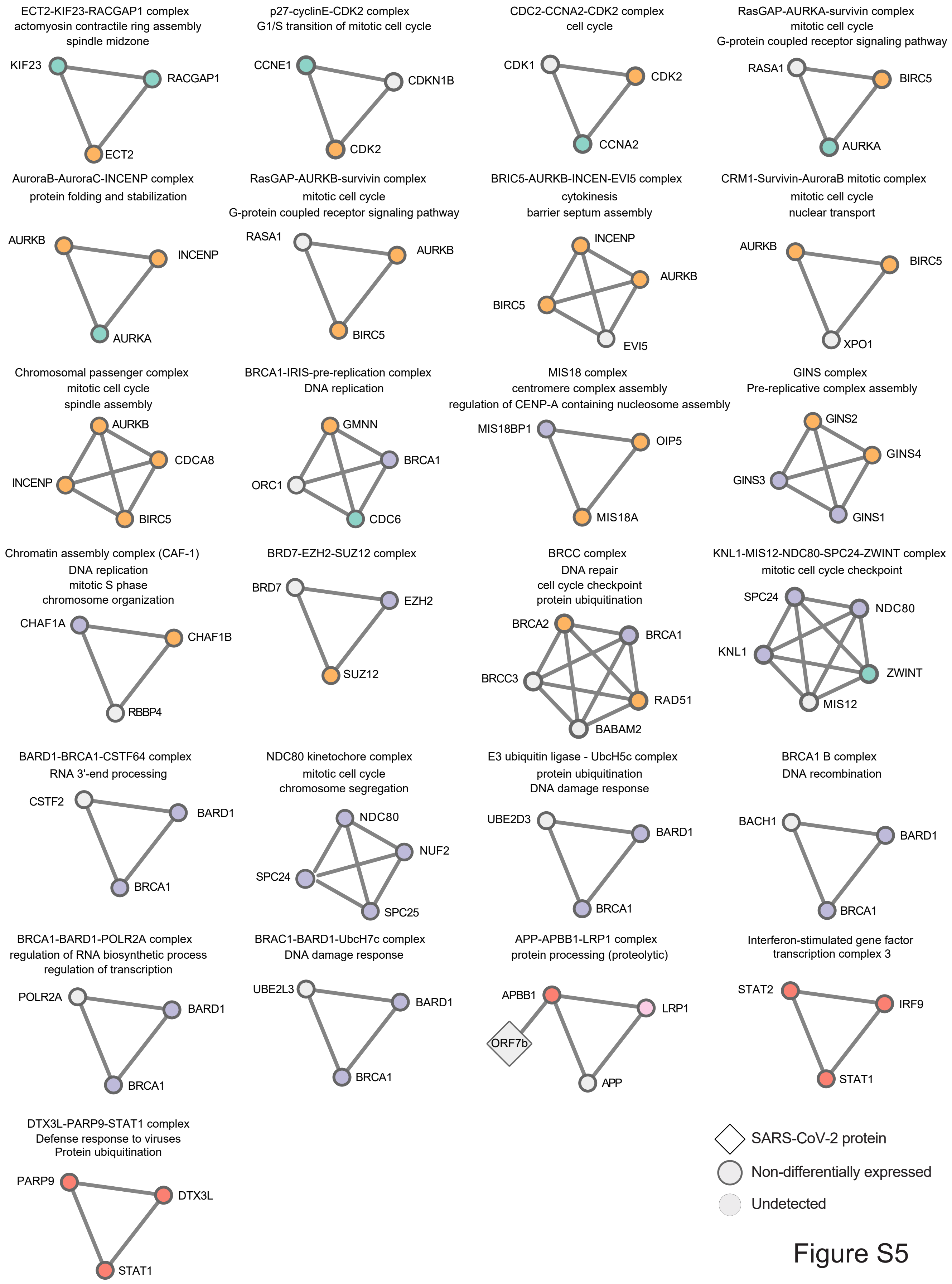

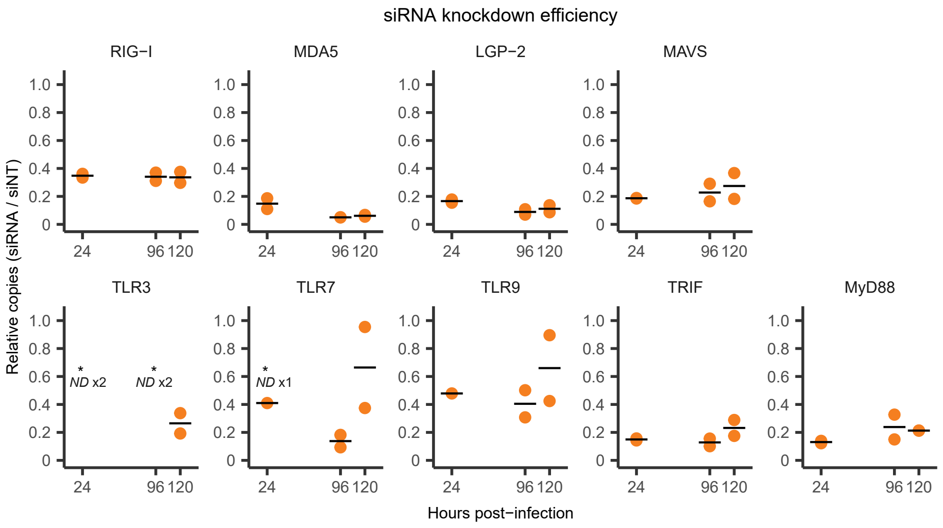
