## Supplemental table S6 for "Systematic analysis of SARS-CoV-2 infection of an ACE2-negative human airway cell"

| REAGENT or RESOURCE | SOURCE | IDENTIFIER |
|---|---|---|
| Oligonucleotides | | |
| SARS-CoV-2 N Forward Primer: 5’-ATGCTGCAATCGTGCTACAA | This paper | N/A |
| SARS-CoV-2 N-Reverse Primer: 5’- GACTGCCGCCTCTGCTC | This paper | N/A |
| SARS-CoV-2 N probe: 5’-FAM/TCAAGGAAC/ZEN/AACATTGCCAA/3IABkFQ | This paper | N/A |
| Human ACE2-Forward Primer: 5’-GGACCCAGGAAATGTTCAGA | This paper | N/A |
| Human ACE2-Reverse Primer: 5’- GGCTGCAGAAAGTGACATGA | This paper | N/A |
| Human TMPRSS2-Forward Primer: 5’- CAGGAGTGTACGGGAATGTGATGGT | This paper | N/A |
| Human TMPRSS2-Reverse Primer: 5’- GATTAGCCGTCTGCCCTCATTTGT | This paper | N/A |
| African green monkey TMPRSS2-Foward Primer: 5’- CAGTCCCAGGCTTCCCTTG | This paper | N/A |
| African green monkey TMPRSS2-Reverse Primer: 5’- TGTTCAGTATGACCTGCCGAG | This paper | N/A |
| RPL13a-Forward Primer: 5’- CATAGGAAGCTGGGAGCAAG | This paper | N/A |
| RPL13a-Reverse Primer: 5’- GCCCTCCAATCAGTCTTCTG | This paper | N/A |
| ACE2-Screen-Forward Primer: 5’-CTCATGATTCATTCATGTCCTTGC | This paper | N/A |
| ACE2-Screen-Reverse Primer: 5’-GACAGTGGGGAAACTTAACTGG | This paper | N/A |
| Human TLR3-Forward Primer: 5’-GGCTAGCAGTCATCCAACAGAA | This paper | N/A |
| Human TLR3-Reverse Primer: 5’- GCAGTCAGCAACTTCATGGC | This paper | N/A |
| Human TLR7-Forward Primer: 5’-GCTGATCTTGGCACCTCTCA | This paper | N/A |
| Human TLR7-Reverse Primer: 5’-TGTCCACATTGGAAACACCATTT | This paper | N/A |
| Human TLR8-Forward Primer: 5’-AGAACAACAGAAACATGGAAAACA | This paper | N/A |
| Human TLR8-Reverse Primer: 5’-TCTTCGGCGCATAACTCACA | This paper | N/A |
| Human TLR9-Forward Primer: 5’-CTTCCCTGTAGCTGCTGTCC | This paper | N/A |
| Human TLR9-Reverse Primer: 5’-TGCGGCAGAAACCCATGCT | This paper | N/A |
| Human MyD88-Forward Primer: 5’-TGATTACCTGCAGAGCAAGG | This paper | N/A |
| Human MyD88-Reverse Primer: 5’-TTCTGATGGGCACCTGGAGA | This paper | N/A |
| Human TRIF-Forward Primer: 5’-CCCGGATCCCTGATCTGCTT | This paper | N/A |
| Human TRIF-Reverse Primer: 5’-GGTGGTGAAGGCATGTTCCA | This paper | N/A |

| | | |
|---|---|---|
| Human RIG-I-Forward Primer: 5’-TGTCCACCTTCAGAAGTGTCT | This paper | N/A |
| Human RIG-I-Reverse Primer: 5’-AGCAGGCAAAGCAAGCTCTA | This paper | N/A |
| Human MDA5-Forward Primer: 5’-AGATGCAACCAGAGAAGATCCA | This paper | N/A |
| Human MDA5-Reverse Primer: 5’-TGGCCCATTGTTCATAGGGT | This paper | N/A |
| Human LGP2-Forward Primer: 5’-TGGGCAAGGCGCAGTTT | This paper | N/A |
| Human LGP2-Reverse Primer: 5’-ACCGAAGCTCCATTCTGCTC | This paper | N/A |
| Human MAVS-Forward Primer: 5’-ACAGCAAGAGACCAGGATCG | This paper | N/A |
| Human MAVS-Reverse Primer: 5’-CGCCGCTGAAGGGTATTGAA | This paper | N/A |
| Human IFIT1-Forward Primer: 5’-CCTCCCTGGAAAATCTAGGCTCT | This paper | N/A |
| Human IFIT1-Reverse Primer: 5’-GTAAAGTGACATCTCAATTGCTCCAGAC | This paper | N/A |
| Human IFIT2-Forward Primer: 5’-AGCTGAGAATTGCACTGCAACCATG | This paper | N/A |
| Human IFIT2-Reverse Primer: 5’-CTCCATCAAGTTCCAGGTGAAATGGC | This paper | N/A |
| Human ISG15-Forward Primer: 5’-CACAGCCCACAGCCCACAG | This paper | N/A |
| Human ISG15-Reverse Primer: 5’-GCTCAGGGACACCTGGAATTCG | This paper | N/A |
| Human MX1-Forward Primer: 5’- CACTGCGAGGAGATCGGTTC | This paper | N/A |
| Human MX1-Reverse Primer: 5’- CTGTTCTCCTGCACCTCCTTG | This paper | N/A |
| Human 18S rRNA-Forward Primer: 5’-CCGCAGCTAGGAATAATGGA | This paper | N/A |
| Human 18S rRNA-Reverse Primer: 5’-CGGTCCAAGAATTTCACCTC | This paper | N/A |
| Probe-V-nCoV2019-S | RNAScope | 848561 |

Table S6, related to STAR methods: Oligonucleotide Sequences
